# The functional genomic circuitry of human glioblastoma stem cells

**DOI:** 10.1101/358432

**Authors:** Graham MacLeod, Danielle A. Bozek, Nishani Rajakulendran, Vernon Monteiro, Moloud Ahmadi, Zachary Steinhart, Michelle M. Kushida, Helen Yu, Fiona J. Coutinho, Ian Restall, Xiaoguang Hao, Traver Hart, H. Artee Luchman, Samuel Weiss, Peter B. Dirks, Stephane Angers

## Abstract

**Summary:** Successful glioblastoma (GBM) therapies have remained elusive due to limitations in understanding mechanisms of growth and survival of the tumorigenic population. Using CRISPR-Cas9 approaches in patient-derived GBM stem cells to interrogate function of the coding genome, we identify diverse actionable pathways responsible for growth that reveal the gene-essential circuitry of GBM stemness. In particular, we describe the Sox developmental transcription factor family; H3K79 methylation by DOT1L; and ufmylation stress responsiveness programs as essential for GBM stemness. Additionally, we find mechanisms of temozolomide resistance and sensitivity that could lead to combination strategies with this standard of care treatment. By reaching beyond static genome analysis of bulk tumors, with a genome wide functional approach, we dive deep into a broad range of biological processes to provide new understanding of GBM growth and treatment resistance.

**Significance:** Glioblastoma (GBM) remains an incurable disease despite an increasingly thorough depth of knowledge of the genomic and epigenomic alterations of bulk tumors. Evidence from multiple approaches support that GBM reflects an aberrant developmental hierarchy, with GBM stem cells (GSCs), fueling tumor growth and invasion. The properties of this tumor subpopulation may also in part explain treatment resistance and disease recurrence. Unfortunately, we still have a limited knowledge of the molecular circuitry of these cells and progress has been slow as we have not been able, until recently, to interrogate function at the genome-wide scale. Here, using parallel genome-wide CRISPR-Cas9 screens, we identify the essential genes for GSC growth. Further, by screening in the presence of low and high dose temozolomide, we identify mechanisms of drug resistance and sensitivity. These functional screens in patient derived cells reveal new aspects of GBM biology and identify a diversity of actionable targets such as genes governing stem cell traits, epigenome regulation and the response to stress stimuli.

## Introduction

Glioblastoma (GBM) is the most common type of primary malignant brain tumor in adults (Wen and Kesari, 2008). Although there are many reasons underlying the treatment-refractory nature of these tumors, inter- and intra-patient tumor heterogeneity and a complete lack of durable responses to therapy are critical contributors to the dismal overall prognosis (Sturm et al., 2014). Temozolomide (TMZ) chemotherapy represents the last major widespread applied advance in GBM therapy (Stupp et al., 2005), although its efficacy is remarkably limited, and it endows tumors with an additional mutational burden that may ultimately drive disease progression (Cancer Genome Atlas Research Network, 2008; Hunter et al., 2006). While large-scale genomic analyses of human GBM tumor samples has revealed a complex genomic architecture (Brennan et al., 2013; Parsons et al., 2008; Uhm, 2009), this has been difficult to link to GBM growth and functional cell properties, particularly due to the tumors’ proliferative and cellular heterogeneity. Indeed, a more comprehensive understanding of molecular determinants of growth and drug responsiveness across this heterogenous cancer is needed, both to identify new therapeutic opportunities and to also provide new strategies that could be partnered with TMZ that is now entrenched in upfront GBM therapy.

GBM growth is thought to be driven by a small subpopulation of GBM stem cells (GSCs) that are capable of self-renewal and generation of progeny that are more limited in their tumorigenic capacity and associated with expression of markers of more differentiated lineages. This concept is supported by recent single cell RNA sequencing approaches (Patel et al., 2014; Venteicher et al., 2017) and our recent fate mapping study of fresh human GBM cells following orthotopic transplantation in mice, which also highlights the presence of a chemotherapy-resistant tumorigenic subpopulation (Lan *et al.,* 2017). Although tumorigenic properties of GBM stem cells are best interrogated *in vivo*, GSC culture systems have enabled expansion of these patient derived cells from many GBM samples for functional experiments on a scale that is difficult to achieve *in vivo* (Pollard *et al.,* 2009). Importantly, these GSC cultures reliably preserve patient specific phenotypic and genotypic characteristics, including at a single clonal level, retain *in vivo* tumorigenic capacity and recapitulate the hierarchical growth behavior observed *in vivo* (Lee *et al.,* 2006; Pollard *et al.,* 2015; Meyer *et al.,* 2015; Lan *et al.,* 2017).

The emergence of the CRISPR-Cas9 technology has enabled genome-wide forward genetic screens, including “cell fitness screens”, which allow for the systematic identification of “core” and “context-specific” essential genes governing cell proliferation across all cell types or in a given cellular genetic background (Aguirre et al., 2016; Shalem et al., 2013; Wang et al., 2013, 2015). Using this approach, we and others reported that approximately 2000 genes are required for the optimal growth of all the cancer cell lines examined, including 400 genes that are unique to each cell type examined (Hart *et al.,* 2015; Wang *et al.,* 2015; Blomen *et al.,* 2015). Context-specific essential genes represent unique genetic vulnerabilities that underlie the concept of synthetic lethality and may, in some cases, be therapeutically exploited. For example, using genome-wide CRISPR-Cas9 essentiality screens, we recently identified the Wnt receptor FZD5 as specifically required for the growth of *RNF43* mutant PDAC cells and showed that FZD5 blocking antibodies robustly inhibit the proliferation of these cells *in vitro* and tumor growth in orthotopic mouse models (Steinhart et al., 2017).

Given the complex patient to patient heterogeneity observed in GBM, additional gene essentiality screens are needed to capture the genetic dependencies common across all patients and essential genes unique to individual GBM genotypes. Here, we performed parallel genome-wide CRISPR screens in eight low passage patient-derived GSC cultures that harbor a range of typical GBM genomic abnormalities matched with their primary tumors (Lan et al., 2017; Pollard et al., 2009) to define the molecular circuitry governing GSC growth and survival. We concurrently performed chemogenomic screens to identify genes modulating TMZ responsiveness revealing mechanisms of therapeutic resistance and strategies for combinatorial therapy. Our results provide the largest series of forward-genetic screens for a solid tumor in patient-derived tumor initiating cells and provide a functional blueprint for GBM tumorigenicity, at the GSC level. These findings identify new GBM vulnerabilities and nominate a diverse repertoire of strategies for additional therapeutic development.

## Results

### Genome-wide CRISPR-Cas9 Screens Identify Genetic Vulnerabilities of GBM Stem Cells

To identify essential fitness genes in GSCs, we performed parallel genome-wide CRISPR-Cas9 screens in 8 well-characterized (Figure S1A) low passage patient-derived GSC cultures engineered to express the Cas9 nuclease (GSC-Cas9) (Figure 1A). Cells were infected with lentivirus carrying the 90K gRNA TKOv1 library (Hart et al., 2015). After selection for infected cells, and T0 sampling, cells were grown for 14 doublings and genomic DNA collected at each passage. We also performed chemogenomic screens (Figure 1A) with lethal or sub-lethal doses (LD90 and LD20 respectively) of TMZ, to identify genes underlying TMZ sensitivity and resistance. The BAGEL algorithm was used to derive a Bayes Factor (BF) for each tested gene by comparing changes in gRNA abundance throughout the course of the screen to experimentally determined gold-standard reference essential and non-essential genes (Hart and Moffat, 2016; Hart et al., 2017). The Bayes Factor is a measure of the confidence that knockout of a specific gene causes a decrease in cell fitness (high BF indicates increased confidence that the knockout of the gene results in a decrease in fitness). In this manner, we identified an average of 1,722 essential genes in each GSC culture at a 5% false discovery rate (FDR) (Table S1). Despite patient to patient genetic differences between these samples, a total of 1,157 genes were hits in at least 5 of the 8 patient-derived GBM cultures screened (Figure S1B). These findings suggest that GSCs require a core set of genes required for proliferation and survival, reminiscent of shared functional properties that we identified in our previous *in vivo* fate mapping study (Lan et al., 2017).

**Figure 1:**
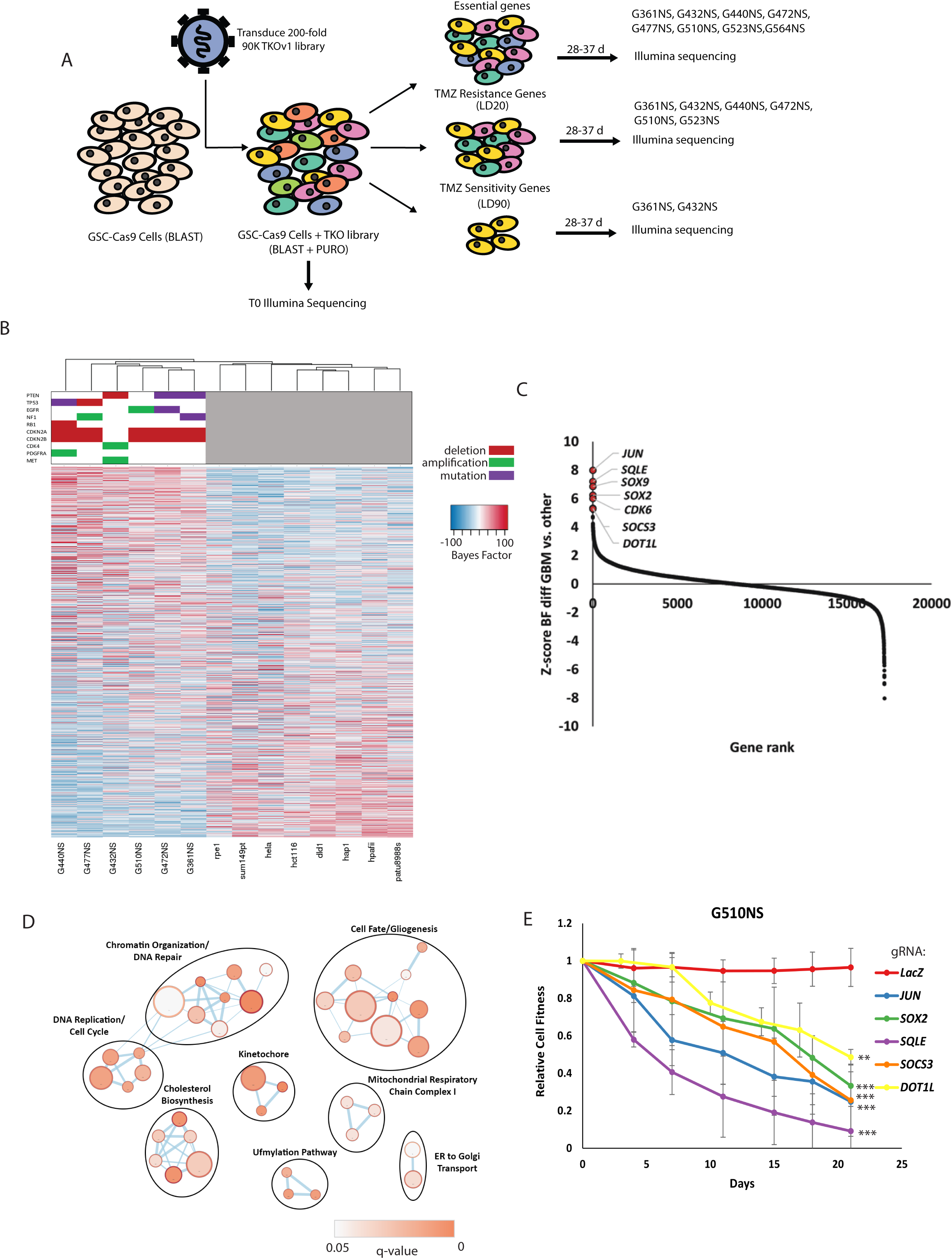
Genome-wide CRISPR-Cas9 screens identify genetic vulnerabilities in patient-derived glioblastoma stem cells. (A) Schematic for genome-wide CRISPR-Cas9 screens for essential genes and modulators of temozolomide sensitivity. (B) Heatmap of normalized gene essentiality Bayes Factor (BF) scores for 1,326 most variable genes in GBM and non-GBM screens. Rank is according to differential essentiality Z scores (Average BF for GSC screens - Average BF for non-GBM screens). GSCs oncoprint for frequently altered genes in GBM is included at the top. (C) Rank order plot depicting differential essentiality between GBM and non GBM screens). Top hits are highlighted. (D) Enrichment Map of GO biological processes and Reactome terms enriched in GBM specific essential genes. Node size represents number of genes belonging to individual terms, colour of node represents significance of the gene set (only q-value < 0.05 gene sets are shown). (E) Multicolor competition assay (MCA) in G510NS-Cas9 showing decrease in cell fitness upon expression of gRNA targeting indicated GBM essential genes (mean ± SD from n=3 replicates, *** p < 0.001, ** p < 0.005 from gRNA-LacZ control). See also Figures S1, S2 and S3 and Tables S1 and S2.

To identify fitness genes that are unique to the stem cell fraction of GBM, we compared the core set of fitness genes derived from our 6 highest quality GSC screens (Figure S1C) to 8 previously reported essentiality screens performed in cancer and epithelial cell lines from diverse tissues of origin (“non-GBM”) (Hart et al., 2015). As expected, GSC screens formed a distinct cluster as defined by the essentiality profile from non-CNS cancer cells, emphasizing the shared context-specific mechanisms governing growth and survival of this cancer type (Figure 1B; Figure S1D). We calculated a Z-score for the difference in average BF-score between GBM and non-GBM screens and derived a ranked list of GSC specific essential genes (Figure 1C; Table S2). Generation of an enrichment map for GBM-specific essential genes using Gene Ontology (GO) Biological Processes and the Reactome knowledge base terms revealed genetic vulnerabilities within a diverse set of networks, including GO terms related to Chromatin Organization and DNA Repair, Cell Fate and Gliogenesis, Cholesterol Biosynthesis, DNA Replication and Cell Cycle, and protein Ufmylation (Figure 1D). To validate genome-wide screening results, we selected 20 conserved GBM-specific essential genes of particular interest on the basis of high Z-score and spanning the key GO processes identified above. We validated all of these hits by testing individual gene knockouts in three GSCs using competitive cell proliferation assays in which we co-cultured GSC-Cas9 cells expressing GFP + a gRNA targeting a gene of interest with cells expressing mCherry + a non-targeting gRNA. The GFP/mCherry ratio was tracked over a period of three weeks (Figure 1E; Figure S2A-F). In most cases, cells expressing the gRNA targeting genes identified in the screen were outcompeted by cells expressing a non-targeting gRNA. We therefore identified a core set of GSC essential genes that represent genetic vulnerabilities crosscutting the majority of GBM patients.

Careful examination of the relationship between gene essentiality profiles and the genetic background of the tested GSCs also identified synthetic lethal interactions unique to patient-specific genetic contexts. For example, 3 of the 8 lines harbored *EGFR* activating mutations or amplification and displayed *EGFR* essentiality, whereas *EGFR* disruption in *EGFR* wild-type lines had no effect on growth (Figure S3A). In addition, given the universal disruption of the Rb pathway in GBM we observed a dependency on the CDK4/6 cyclin-dependent kinase family (Wiedemeyer et al., 2010). We saw a specific reliance on *CDK4* or *CDK6* depending on whether the cells had *CDK4* amplification or *CDNK2A/B* disruption (Figure S3B). We validated these findings using single gene knockouts of *CDK4* and *CDK6* in 5 GSC cultures, including one not included in the screen panel. Consistent with our screen results, the proliferation of GSCs harboring *CDK4* amplification was exclusively dependent on *CDK4*, while GSCs with *CDKN2A/B* deletion were exclusively dependent upon *CDK6* (Figure S3C). These findings illustrate the potential for genome-wide CRISPR-Cas9 functional genomics screens to identify both core and genotype-specific therapeutic vulnerabilities in the GBM patient population. In turn, this provides exciting insights into novel therapeutic strategies for this devastating disease.

### Genetic Programs Governing Stemness are Essential in GSC Cultures

One striking GSC essential GO network contained genes important for cell fate and gliogenesis. Importantly, we identified genes associated with stem cells such as members of the Sox family of transcription factors, *OLIG2* and *SALL1* (Figure 2A). These findings are consistent with previous studies demonstrating a role for these genes in normal neural precursor compartments and their capacity to reprogram more differentiated GBM cells into a stem cell state and sustain tumor propagation *in vivo* (Suvà et al., 2014).

**Figure 2:**
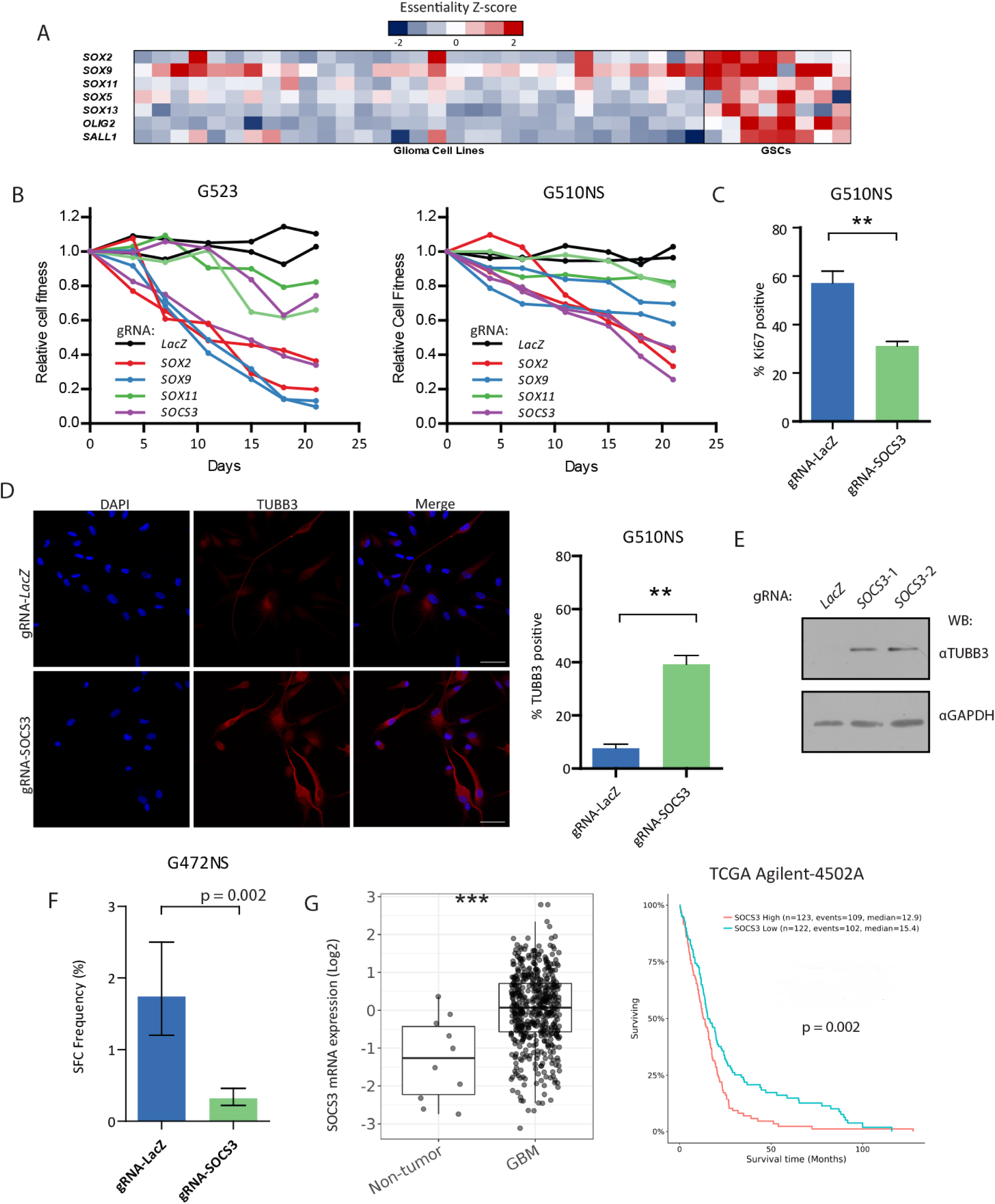
Essential regulators of GSC fate identified in genome-wide CRISPR-Cas9 screens. (A) Heatmap of normalized gene essentiality for neurodevelopmental transcription factors governing stem cell state in GSCs. (B) Multi-colour competition growth assays (MCA) for GSC expressing the indicated gRNAs targeting Sox family genes and *SOCS3* in two GSC cultures. Relative cell fitness is expressed as mean ± SD from n=3 replicates. (C) Quantification of proliferating cells using Ki67 staining in G510NS-Cas9 cells infected with *LacZ* control or *SOCS3* gRNAs. Error bars, mean ± SEM from n=3 biological replicates. (D) Left: Representative Immunofluorescence staining of TUBB3 in G510NS-Cas9 cells infected with gRNA-LacZ or gRNA-SOCS3. Nuclei were stained with DAPI. Scale, 50 µm. Rights: quantification of TUBB3+ cells. Error bars, mean ± SEM from n=3 biological replicates. (E) Western blotting of TUBB3 in G510NS-Cas9 cells infected with *LacZ* control or *SOCS3* gRNAs. GAPDH loading control included. Images are representative of 3 biological replicates. (F) Sphere initiating frequency of G472NS-Cas9 GSCs following expression of gRNAs for *LacZ* or *SOCS3*. Data represents mean ± upper and lower 95% confidence intervals. (G) Left: *SOCS3* mRNA expression in TCGA Glioblastoma dataset. Right: survival analysis for GBM patients in TCGA dataset segregated by high and low *SOCS3* expression. ** p-value < 0.005, *** p=value < 0.001

In particular, *SOX2* and *SOX9* were found to be essential in 5 and 6 cultures respectively, supporting their role as core regulators of stemness in GSCs. The GSC screens also identified *SOX13*, *SOX11* and *SOX5* as differentially essential across GSC cultures suggesting patient specific stem cell transcription factor programs that cooperatively sustain GSC self-renewal in patient-specific contexts. Individual knockout of *SOX2* and *SOX9* in cell competition assays confirmed strong negative effects on cell proliferation whereas *SOX11* disruption conferred a milder and more variable anti-proliferation phenotype (Figure 2B; Figure S2C-E). We conclude that stem cell gene networks fuel the growth of patient-derived GSCs. Strikingly, comparison of our screens with recently reported CRISPR-Cas9 screens, performed in 31 serum-based glioma cell lines (Meyers et al., 2017) and two GSCs (Toledo et al., 2015) showed a different repertoire of essential genes (Figure S3D). Notably, the cell fate and gliogenesis genes were unique to our screens, perhaps reflecting the different growth conditions and duration of analysis of dropouts (Figure 2A). Indeed, while GSCs cultured in growth factor supplemented stem cell media have been demonstrated to be a robust model of GBM and are highly enriched for tumor initiating cells *in vivo*, serum cultured glioma cell lines are less capable of tumor formation and bear little histological resemblance to patient disease (Lee et al., 2006; Pollard et al., 2009).

Taking advantage of this unique feature, we next asked whether additional, less well recognized stem cell genes could also be identified. Among the top scoring GBM-specific essential genes was Suppressor of Cytokine Signaling 3 (*SOCS3*), which has been previously shown to maintain stemness in neural stem cells (Cao et al., 2006). Intriguingly, *SOCS3* was reported to have opposing effects in glioma and function as either a tumor suppressor or oncogene depending on context (Jiang et al., 2017; Martini et al., 2008; Zhou et al., 2007). To gain a better understanding of the functional mechanisms underlying *SOCS3* in GSCs, we first knocked out its expression with 2 individual gRNAs to validate its essentiality in multiple GSC cultures. We found that *SOCS3* knockout led to a reduction in GSC proliferation as measured by competitive cell proliferation assays (Figure 2B; Figure S2B-E) and Ki67 staining (Figure 2C), increased expression of the neuronal marker TUBB3 (Figure 2D and 2E) and decreased sphere forming ability in limiting dilution assays (Figure 2F). Analysis of TCGA datasets revealed an upregulation of *SOCS3* levels in GBM when compared with non-tumor tissue and an association between high *SOCS3* expression and worse prognosis (Figure 2G). Together these results imply a role of SOCS3 in GSC stemness and underlie its importance during GBM progression.

### The Histone Methyltransferase DOT1L Regulates Stemness and Proliferation in GSCs

Our previous GBM fate mapping work pointed to epigenetic compounds as disruptors of GSC growth (Gallo et al., 2015; Lan et al., 2017). In line with these findings, our screens identified epigenetic dependencies that are unique to GSCs compared to other non-GBM cancer types, suggesting an opportunity to develop CNS specific epigenetic modifiers for GBM treatment. We observed the epigenetic modulators *DOT1L*, *EZH2* and *SUV420H1* were among our top scoring hits as GSC-specific essential genes. The H3K79 methyltransferase DOT1L is highly expressed in GSCs compared to bulk tumor (Figure 3A) and is an essential gene in 7 of 8 GSC cultures, supporting a near pan-GSC essential function. We next validated *DOT1L* essentiality in GSCs using CRISPR-Cas9 gene editing (Figure 1E; Figure S2C,D and F) and found that the clinically relevant small molecule inhibitor of DOT1L, EPZ-5676, inhibited GSC growth (Figure 3B). The effects of EPZ-5676 were prominent but only after longer periods, consistent with inhibition of an epigenetic process (Figure 3B,G; Figure S5A, B). Importantly, in line with regulation of a stem cell program, morphologic changes consistent with differentiation were observed in a sphere culture model of GSCs (Figure S5A) with increased expression of neuronal and glial markers and decreased expression of stem cell markers (Figure 3C and D; Figure S4A,B and C) after 7 days of treatment. These changes translated into marked reduction in self-renewal as measured by assays of sphere forming capacity and, in addition, reduction of cellular migration/invasion (Figure 3E and F; Figure S4D and E). Critically, assessment of histone marks following treatment of GSCs with EPZ-5676 demonstrated on-target DOT1L inhibition as H3K79me2 levels were specifically decreased without affecting other histone marks (Figure S4F), and this was further diminished over time (Figure S5B). Consistent with this observation, we found that longer treatment periods of 21 days led to decreased BrdU incorporation, G0/G1 cell cycle arrest, as well as increased apoptosis (Figure 3B and G; Figure S5C). Importantly, EPZ-5676 treatment led to a significant loss of the H3K79me2 mark at the key neural stem cell regulators and GBM-specific essential genes *SOX2* and *OLIG2* (Figure 3H). Therefore, GSC stemness properties are attenuated through inhibition of H3K79me2.

**Figure 3:**
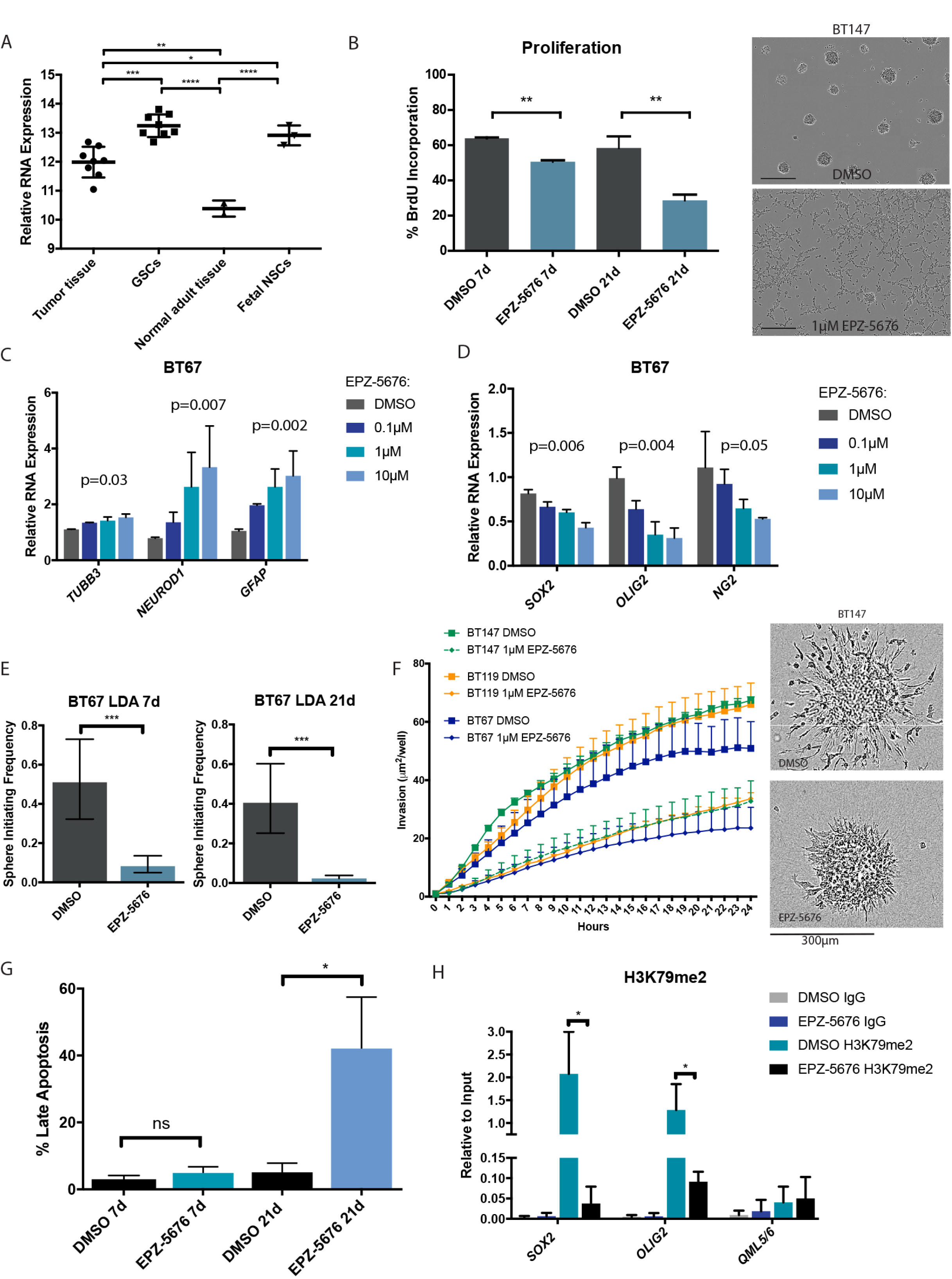
DOT1L epigenetically regulates GSC stem cell properties. (A) Expression of *DOT1L* in GBM primary tumor tissue, GSCs, normal adult brain tissue and fetal neural stem cells (NSCs) (one-way ANOVA, * p < 0.05, ** p <0.005, *** p 0.0001, **** p < 0.0001). (B) The effect of EPZ-5676 treatment on GSC proliferation (left, unpaired t-test ** p < 0.005, data are represented as mean ± S.D). (C) Gene expression of neuronal differentiation markers (*TUBB3* and *NEUROD1*) and astrocytic differentiation marker (*GFAP*) in BT67 GSCs treated with EPZ-5676. Data are represented as mean ± S.D. (D) Gene expression of stem and progenitor cell markers (*SOX2*, *OLIG2* and *NG2*) in BT67 GSCs treated with EPZ-5676. Data are represented as mean ± S.D. (E) Sphere initiating frequency of BT67 GSCs following short-term and long-term EPZ-5676 treatment. Data represents mean ± upper and lower 95% confidence intervals. (F) Quantified invasion of GSC lines following EPZ-5676 pre-treatment (left) representative image depicting invasion of GSC culture BT147 (right) (Two-way ANOVA p-value BT67=0.02, p-value BT119=0.002, p-value BT147=0.0006). Data are represented as mean ± S.D. (G) Percentage of late apoptotic GSCs treated short-term and long-term with EPZ-5676 (unpaired t-test * p <0.05). Data are represented as mean ± S.D. (H) Enrichment of H3K79me2 levels within *SOX2* and *OLIG2* gene regions as well as negative control heterochromatin region *QML5/6* following EPZ-5676 treatment (unpaired t-test * p <0.05). See also Figures S4 and S5.

To investigate the potential of targeting DOT1L for the treatment of GBM, we tested the effect of EPZ-5676 treatment on GSCs in an immunodeficient mouse model. As our data support that EPZ-5676 poorly crosses the blood brain barrier (Waters et al., 2016) in control animals (Figure 4A), we pretreated GSC cultures *in vitro* followed by orthotopic transplantation. Animals that received tumors pre-treated with EPZ-5676 had reduced tumor growth and longer survival (Figure 4B, C and D; Figure S5D,E and F). Importantly, dosing of mice with EPZ-5676 led to tumor growth inhibition following subcutaneous establishment of tumors. In these *in vivo*-treated tumors, we saw erasure of the H3K79me2 mark and increased expression of genes associated with differentiation (Figure 4E, F). Given these data, further pre-clinical studies are warranted to investigate DOT1L histone methylation in GBM, particularly if more CNS penetrant compounds can be developed. The efficacy across multiple patient-derived cell populations supports broad targeting of processes such as chromatin regulation (Bulstrode et al., 2017; Jin et al., 2017; Lan et al., 2017), which could be a strategy to treat the diversity of GBM genotypes, and overcome GBM interpatient heterogeneity.

**Figure 4:**
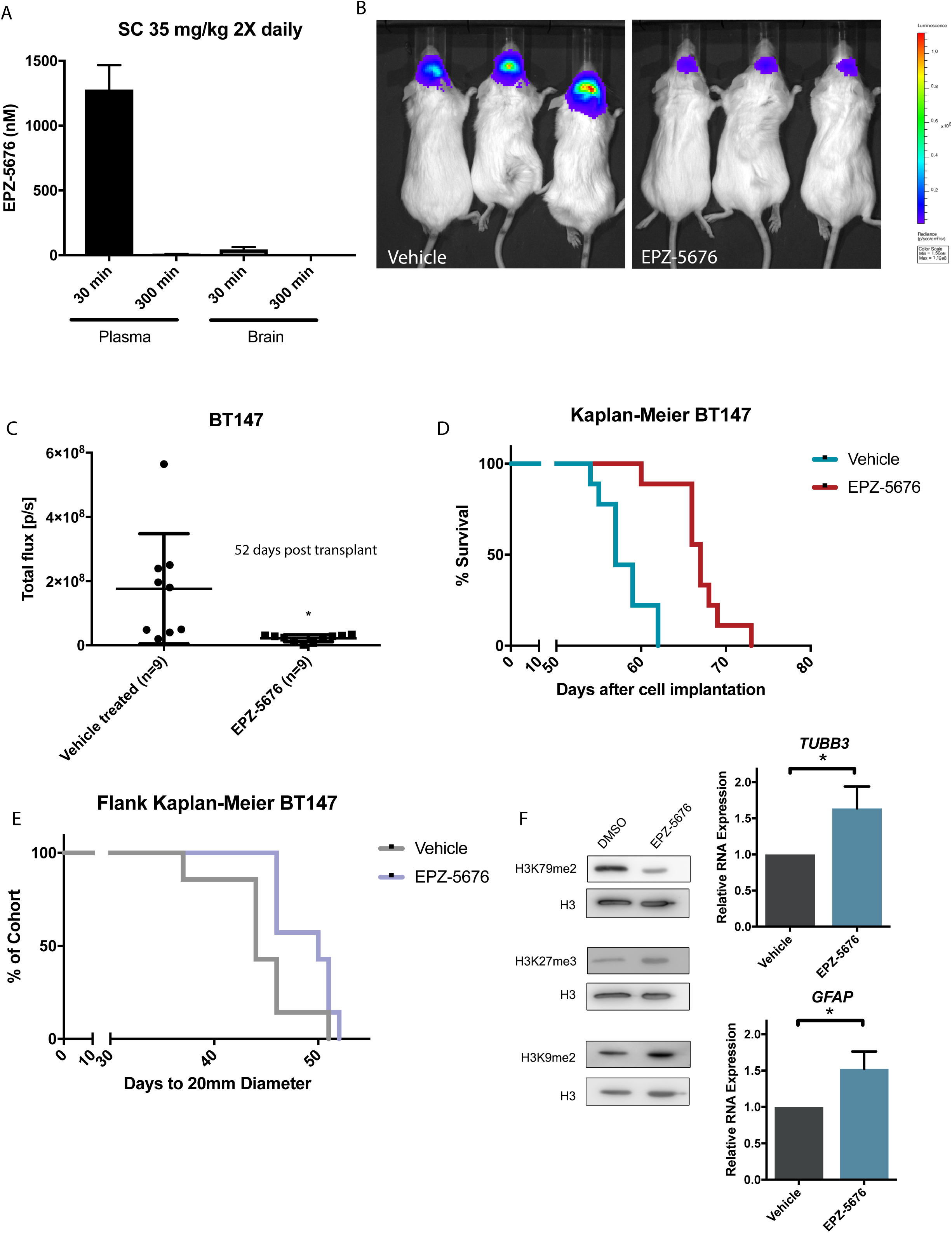
EPZ-5676 treatment decreases GBM tumor growth and improves overall survival. (A) Analysis of EPZ-5676 penetration in the brain following 30 minutes and 5 hours of treatment administration by subcutaneous (SC) injection. Data represented as mean ± S.D. (B) Representative tumor burden at 52 days post engraftment for mice that were orthotopically xenografted with luciferase labelled BT147 that were pre-treated with vehicle or EPZ-5676 for 28 days. (C) Quantified tumor burden for mice that were orthotopically xenografted with BT147 that were pre-treated with vehicle or EPZ-5676 for 28 days (unpaired t-test * p < 0.05). (D) Kaplan-Meier survival curves for EPZ-5676 pre-treated BT147 in an orthotopic model (Mantel-Cox test **** p < 0.0001). (E) Kaplan-Meier survival analysis following subcutaneous EPZ-5676 treatment in a flank xenograft model (Mantel-Cox test * p < 0.05). (F) H3K79me2 as well as control histone marks H3K27me3 and H3K9me2 in flank tumor samples collected from vehicle or EPZ-5676 treated mice (left). Gene expression of neuronal (*TUBB3*) and astrocytic (*GFAP*) markers in flank tumors (unpaired t-test * p < 0.05). See also Figure S5.

### Regulators of Cell Stress Response Pathways are Essential in GSCs

Several highly ranked GBM-specific essential genes were also enriched with functions within stress response pathways. For example, *c-JUN* was the top ranked GBM-specific essential gene across our panel of GSC cultures, and the upstream kinase *MAP2K7* was also identified (Figure 5A). The JNK-JUN pathway functions to protect cells from stress-induced apoptosis (Dhanasekaran and Reddy, 2008) and has recently been reported to be activated in GBM (Matsuda et al., 2012; Yoon et al., 2012). Individual CRISPR-Cas9 mediated knockout of either c-*JUN* or *MAP2K7* significantly reduced cell fitness in multiple GSC cultures (Figure 1A; Figure S2C-E), validating the screen results. Additionally, treatment of GSC cultures with the pan-JNK inhibitors SP600125 and bentamapimod caused a robust reduction in cell viability (Figure 5B).

**Figure 5:**
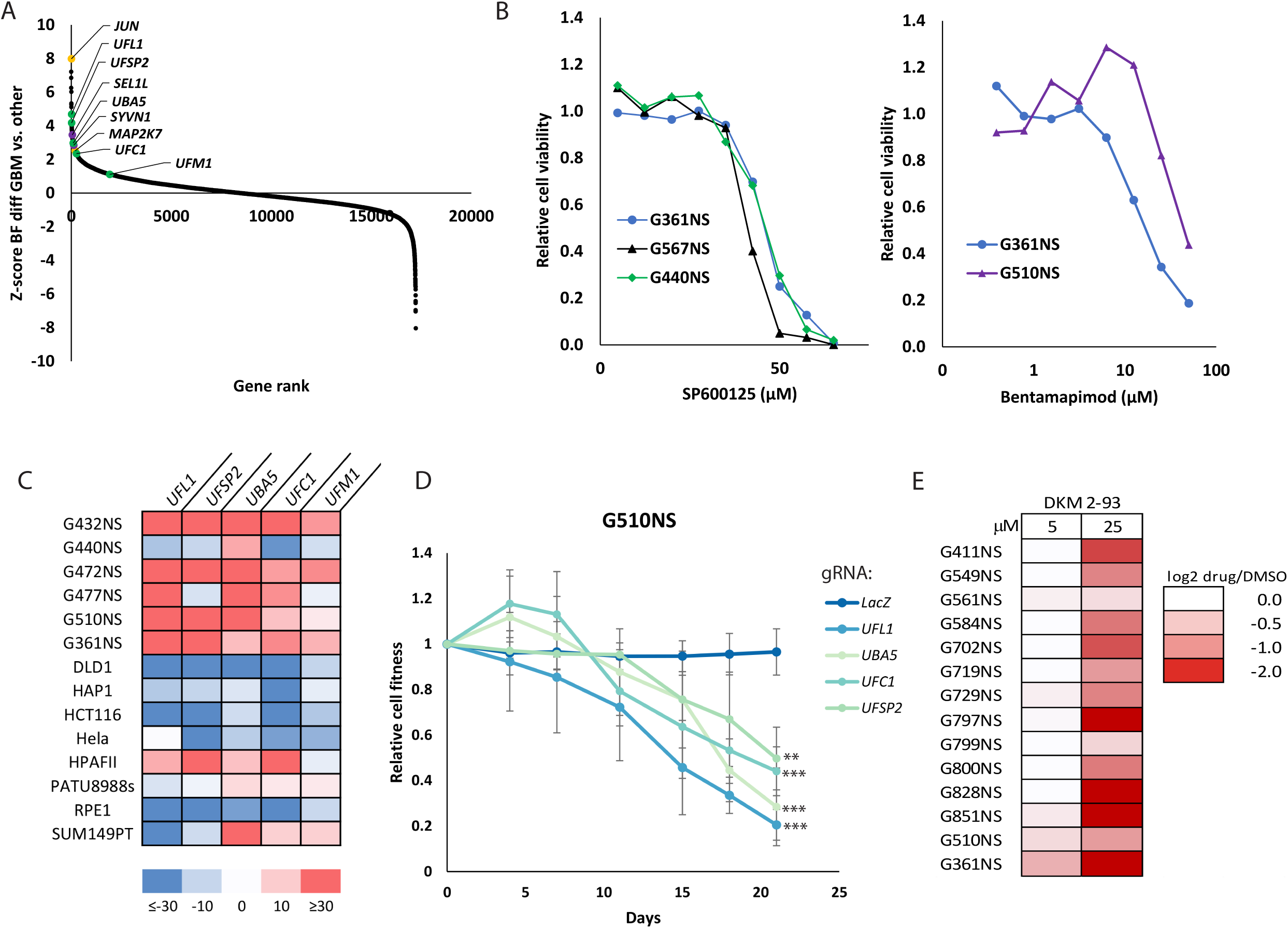
Essential Regulators of Cell Stress in GSCs. (A) Rank order plot depicting differential essentiality between GBM and non GBM screens). Stress signaling pathway components are highlighted: JNK-Jun signaling (yellow), ERAD (purple) and Ufmylation (green). (B) Dose-response curves performed with two JNK inhibitors, SP600125 (left) and Bentamapimod (right), in GSCs, n=3 biological replicates. (C) Heatmap of normalized gene essentiality BF scores for members of the Ufmylation pathway in GBM and non-GBM screens. (D) Multi-colour competition growth assays (MCA) for GSC expressing the indicated gRNAs targeting Ufmylation pathway genes. Relative cell fitness is expressed as mean ± SD from n=3 replicates, *** p < 0.001, ** p < 0.005 from gRNA-LacZ control. (E) Relative cell viability of GSCs treated with UBA5 inhibitor DKM 2-93.

One recently uncovered stress-related pathway overrepresented amongst GBM-specific essential genes was the protein ufmylation system (Figure 1D), a ubiquitin-like post-translational modification linked to ER stress during development and stem cell homeostasis (DeJesus et al., 2016; Wei and Xu, 2016; Zhang et al., 2012). All known ufmylation pathway members were striking hits across the screened GSCs (Figure 5C), suggesting that this pathway may be highly critical for the maintenance of the stem cell compartment in GBM. To further test this dependency on the ufmylation system, we first knocked out the four highest ranking genes in this pathway and subjected these cells to competitive cell growth assays with control cells. Single gene knockout of *UBA5* (E1), *UFC1* (E2), *UFL1* (E3) and *UFSP2* (UFM1 protease) all caused a significant reduction of cell fitness (Figure 5D) confirming that disruption of any of the key catalytic steps in this pathway represents a potential therapeutic vulnerability. Next, we treated a panel of 14 patient-derived GSCs with DKM 2-93 (Roberts et al., 2017), a small molecule inhibitor of UBA5. We observed a robust inhibition of proliferation (Figure 5E) confirming the essentiality of this pathway across all tested GSC cultures. The Endoplasmic Reticulum-Associated Degradation (ERAD) pathway (Kim et al., 2015) members *SEL1L* and *HRD1* (*SYVN1*) were also identified as GBM specific essential genes (Figure 5A). We conclude that GSCs are under high proteotoxic stress and as a result rely on activation of different stress signaling pathways for survival. These results also hint at an opportunity to target the ufymylation proteostasis gene network as a potential therapeutic strategy for GBM.

### Chemogenomic screens reveal mechanisms of Temozolomide sensitivity in GSCs

The GBM chemotherapy agent temozolomide has poor efficacy and concerns exist with its mutagenesis propensity due to DNA alkylation. However, it is used ubiquitously given that a small subset of patients displays longer term survival. For patients who derive no benefit from TMZ treatment, it is necessary to nominate potential treatment combinations that may expand clinical efficacy for this current chemotherapy standard of care. To achieve this goal, further studies are needed to understand the genetic mechanisms of intrinsic TMZ resistance. We therefore performed additional CRISPR-Cas9 screens in the context of low dose and high dose TMZ (Figure 1A). Positive selection screens were performed in two patient-derived GBM cultures (G361NS, G432NS) treated with a lethal dose of TMZ (100 µM) to define genes required for TMZ sensitivity. This dose of TMZ resulted in over 90% cell death over 10 days of treatment, with resistant clones emerging thereafter. Sequencing of integrated gRNAs in resistant cells after 3 weeks of TMZ treatment revealed dramatic enrichment for gRNAs targeting only four genes, the core members of the mismatch repair (MMR) pathway *MLH1, MSH2, MSH6* and *PMS2* (Figure 6A, Figure S6A). The MMR pathway initiates futile cycling of attempted DNA repair in response to TMZ lesions and is required for its cytotoxicity (Wang and Edelmann, 2006). Since *MSH6* mutations are commonly found in recurrent GBM (Hunter et al., 2006; Lan et al., 2017), the screen hence faithfully models tumor evolution *in vitro.* Validating these results, introduction of *MLH1* and *MSH2* mutations in GSCs conferred acquired TMZ resistance (Figure 6B). Given that the four core MMR genes were the sole hits in two separate genome-wide screens, we conclude that this pathway is the primary mediator of TMZ sensitivity, and that further clinical efforts can be directly focused on MMR integrity to improve treatment responsiveness.

**Figure 6:**
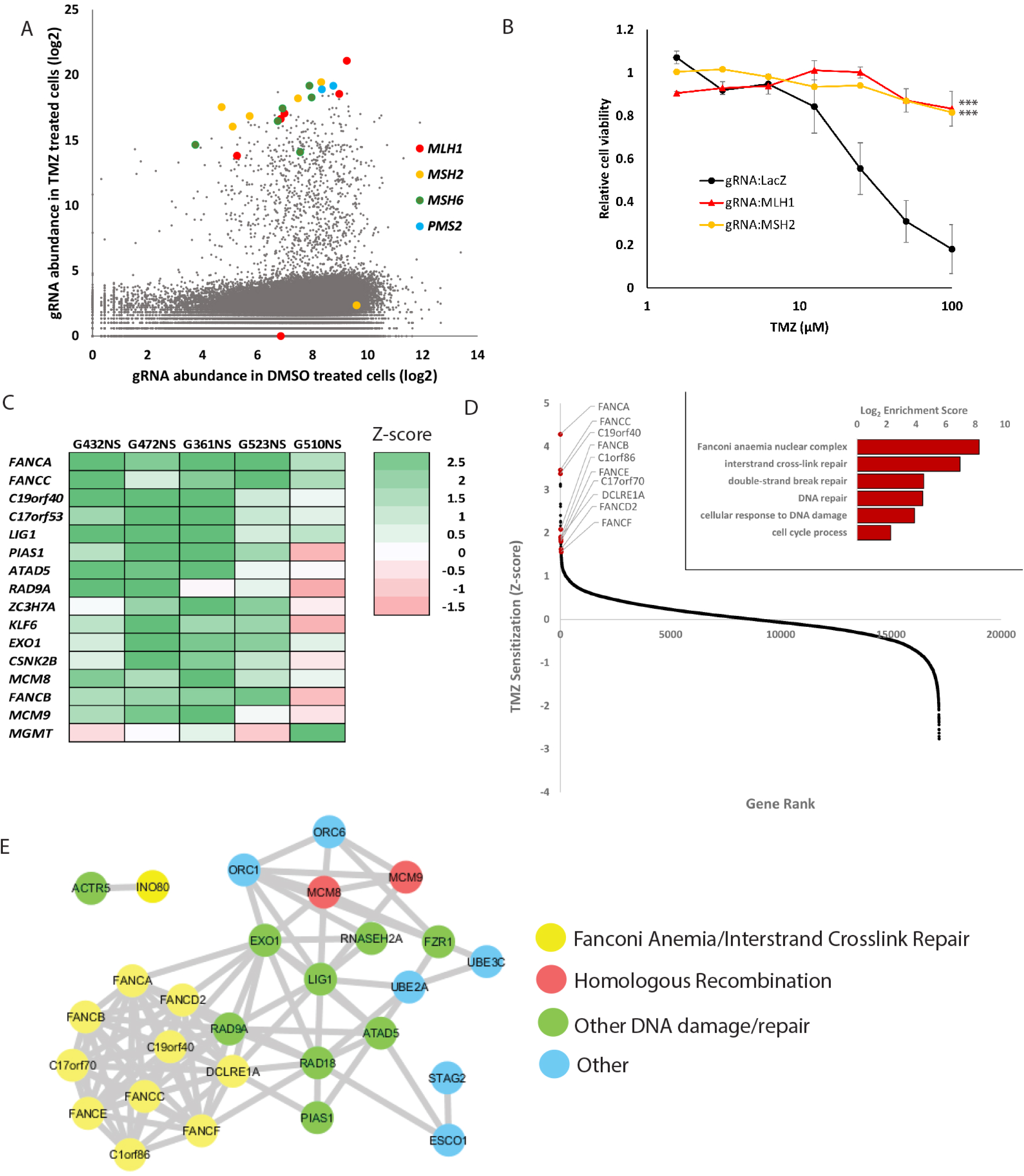
Chemogenomic screens identify modulators of temozolomide sensitivity in GBM. (A) Scatterplot of sequencing reads for gRNAs obtained in TMZ sensitivity screen (LD90, positive selection) with enriched MMR pathway genes highlighted. (B) TMZ dose response curve for GSCs harbouring indicated MMR pathway gene knockout. Mean ± SEM for n=3 biological replicates, *** = p < 0.001. (C) Results of TMZ resistance screens represented as Z-scores of differential BF values obtained for TMZ (LD20, negative selection) and vehicle treated GSCs. Positive Z-scores indicate gene disruption confers TMZ hypersensitivity. (D) Rank order plot indicating genes implicated in TMZ intrinsic resistance, top hits include multiple members of the Fanconi Anemia and Interstrand crosslink repair pathways. Inset shows GO terms enriched. (E) STRING network analysis of top 50 genes identified in TMZ resistance screens showing significant PPI enrichment score (p-value < 1.0e-16). See also Figure S6 and Tables S3 and S4.

To identify mechanisms of intrinsic resistance of the GSC population to TMZ, we performed negative selection screens in the presence of a sublethal dose of TMZ (LD20) in 5 patient-derived GSCs (Figure 1A). Z-scores reflecting the difference in gene essentiality in the presence or absence of TMZ revealed that 4/5 cultures displayed a core subset of genes amongst the top TMZ sensitizers (Figure 6C; Table S3). GO term enrichment (Figure 6D and Table S4) and STRING analysis (Szklarczyk et al., 2017) revealed gene and protein networks involved in multiple DNA repair pathways. Notably, GO terms for the Fanconi Anemia/interstrand crosslink repair and homologous recombination pathways were the most significantly enriched (Figure 6D and 6E). Supporting these results, individual knockout of the Fanconi Anemia pathway genes *FANCA* and *C19orf40* confirmed sensitization of GSCs to TMZ (Figure S7A). Conversely, analysis of the TMZ-resistant G510NS revealed the TMZ adduct repairing enzyme *MGMT* as the top hit restoring TMZ sensitivity (Figure 6C), consistent with RNA-seq results showing this line to be the only *MGMT* expressing line in our screen panel. The identification of a core set of genes involved in intrinsic TMZ resistance across multiple, heterogeneous GSCs, many of which with potentially druggable enzymatic functions, suggests rational strategies for the development of combinatorial therapies. This further provides novel strategies for targeting mechanisms of intrinsic resistance to hypersensitize tumor cells to standard of care chemotherapy.

Along these lines, *MCM8* and *MCM9* which together form a dimeric helicase complex involved in homologous recombination, downstream of the Fanconi Anemia/interstrand crosslink repair pathways as well as the DNA mismatch repair pathway (Nishimura et al., 2012; Traver et al., 2015), were top hits in multiple TMZ chemogenomic screens. We performed functional experiments to determine the effects of these genes on TMZ responsiveness. Knockout of *MCM8* or *MCM9* had no significant effect on cell fitness in untreated cells but strongly sensitized GSC lines to TMZ and had little effect in human neural stem cells (Figure 7A and 7B; Figure S7A-D). Treatment of GSCs with low dose TMZ resulted in double strand breaks that were rapidly repaired in cells expressing a non-targeting gRNA, while cells expressing gRNAs targeting *MCM8* and *MCM9* displayed persistent γH2AX foci, a hallmark of DNA repair defects (Figure 7C and 7D). We further validated the role of MCM8 in contributing to TMZ resistance using a Dox-inducible *MCM8* construct, which rescued the TMZ hypersensitivity displayed by *MCM8* knockout GSCs (Figure S7E). It has been previously shown that the cancer stem cell population demonstrates a highly efficient DNA damage response system which contributes to disease recurrence (Bao et al., 2006). Given their enzymatic function and previous success in development of small molecule DNA helicases inhibitors, we nominate the MCM8/9 complex as a target to be addressed in the context of TMZ. This finding reinforces the power of chemogenomic screens to identify mechanisms of intrinsic drug resistance that could be therapeutically leveraged for more efficient treatments.

**Figure 7:**
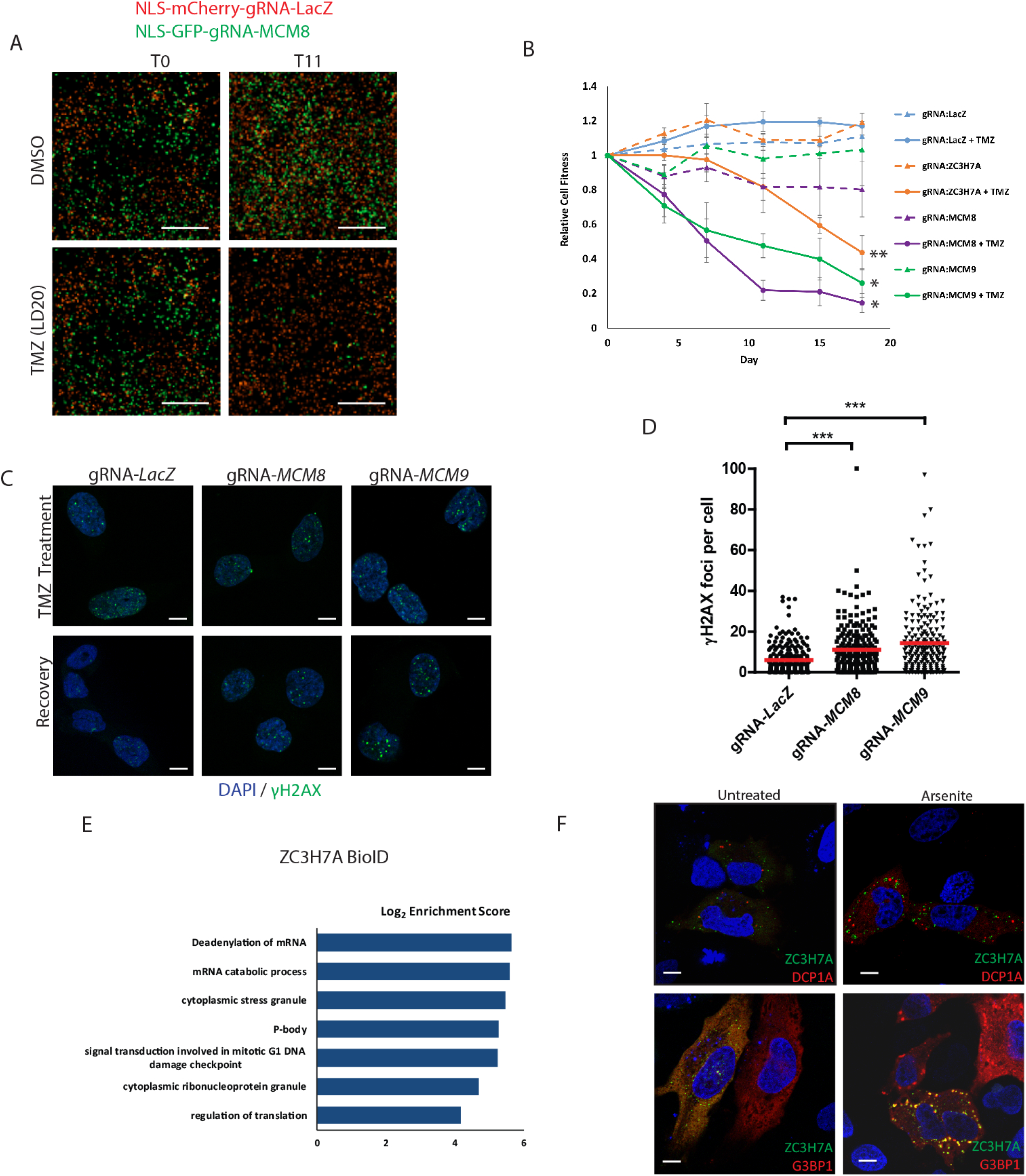
Genes involved in TMZ intrinsic resistance. (A) Representative multi-colour competition assay image showing hypersensitivity to TMZ when the intrinsic resistance gene *MCM8* is knocked out. Note the decrease in green cells (expressing gRNA targeting *MCM8*) between T0 and T11 in the presence of low dose (LD20) TMZ. Scale, 1,000 µm. (B) G361NS-Cas9 MCA results for n=3 replicates, mean ± SEM indicate knockout of *MCM8*, *MCM9* and *ZC3H7A* confer TMZ hypersensitivity. (C) Delayed repair of TMZ induced double-stranded break in *MCM8* and *MCM9* knockout GSCs as visualized by γH2AX immunofluorescence. Top: GSC nuclei at 24 hours TMZ treatment, bottom: 72 hours post-treatment. Scale, 10 µM (D) Quantification of γH2AX foci after 72 hours recovery, from n=3 biological replicates, *** indicates p-value < .0001, Fisher’s Exact Test (E) GO Biological Process and Cellular Component enrichment in proteins identified by BioID of ZC3H7A. (F) Venus-ZC3H7A localizes to cytoplasmic granules and co-localizes with the stress granules marker G3BP1 but not the P-body marker DCP1A when cells are stressed with sodium arsenite. Scale, 10 µM. See also Figure S7 and Tables S5 and S6.

A number of completely uncharacterized genes were also identified to mediate intrinsic TMZ resistance in GSCs. *C17orf53* and *ZC3H7A* were each identified as hits in at least 3 GSC cultures (Figure 6C). Knockout or overexpression of the zinc finger gene *ZC3H7A* led to increased sensitivity and resistance to TMZ respectively (Figure 7B; Figure S7F). To determine the function of ZC3H7A, we performed BioID affinity purification-mass spectrometry to identify ZC3H7A-associated proteins (Table S5). Proximal interactors of ZC3H7A were enriched for biological processes relating to regulation of translation, mRNA processing, cytoplasmic stress granule and p53-mediated DNA damage signaling (Figure 7E; Figure S7G, Table S6). Expression of Venus-tagged full-length ZC3H7A cDNA in both Hela and GSCs showed localization in cytoplasmic granules (Figure 7F, Figure S7H). Since known stress granule (UBAP2L, G3BP1) and RNA Processing body (CNOT complex) proteins were found to interact with ZC3H7A in our BioID experiments, we tested whether ZC3H7A colocalized with markers of either structure. We found that ZC3H7A localized to cytoplasmic granules that co-localized with the stress granule marker G3BP1 upon sodium arsenite treatment (Figure 7F, Figure S7H and I), but did not significantly co-localize with the P-body marker DCP1A. ZC3H7A is therefore a novel stress-responsive regulator of chemoresistance in GBM.

## Discussion

There has been disappointingly slow progress in identifying effective therapies for GBM, in part due to a lack of understanding of the genes essential for functional properties of these tumors and of GSCs, the most primitive cell types shown to lie at the root of GBM growth and resistance (Chen et al., 2012; Lan et al., 2017). The complex cellular ecosystem of GBM tumors certainly suggests that we still have much to learn about cellular mechanisms of tumor growth, particularly *in vivo*, but we need accessible human cell models to interrogate functional properties of enriched populations of tumorigenic cells. Fortunately, patient-derived GSC cultures represent a relatively simple system to generate a cell model of GBM on a very efficient patient to patient basis. These cells, as close as currently attainable, reflect the properties of primary GBM tumors (Lan et al., 2017; Lee et al., 2006; Meyer et al., 2015; Pollard et al., 2009) including following orthotopic xenograftment (Ben-David et al., 2017; Lan et al., 2017). Critically, these GSC cultures are scalable for functional interrogation at the genome-wide level.

In this study, parallel genome-wide CRISPR-Cas9 screens in multiple patient-derived GSCs, identified core growth programs for this heterogeneous disease and uncovered mechanisms underlying TMZ resistance and sensitization. The identification of common GSC essential genes, despite inherent intratumoral heterogeneity and irrespective of patient tumor genotype, provides insight into the underlying biology of GBM and identifies new avenues for drug therapy. In particular, it suggests genes and pathways that are important for GSC properties that were not previously identified following genomic characterization. Moreover, our findings point to a convergence of the large diversity of GBM molecular changes on a potentially smaller number of essential pathways, likely based on aberrant stemness function. Comparison of our GSC screens to our previously performed essentiality screens in cancer and epithelial cell lines from diverse tissues of origin also identifies genetic vulnerabilities unique to GSCs, suggesting specific targeting strategies. These findings provide evidence of the GBM-specific mechanisms used to maintain cancer stem cell characteristics and promote GSC survival.

Equally important is the combination of functional approaches with a detailed understanding of the GBM mutational landscape to begin linking genotypes with phenotypes. By performing a limited number of genome-wide CRISPR essentiality screens in GSCs, characterized by exome-sequencing and RNA-seq, we were able to identify genotype-specific essential genes. For example, within the frequently altered Retinoblastoma pathway, we identified genotype-specific dependencies on either *CDK4* or *CDK6*. GSCs with amplification of *CDK4* were predictably reliant on *CDK4* for survival. However, we also found that GSCs with *CDKN2A/B* deletion had an unexpected dependency on *CDK6* alone, despite sharing common regulation and function with *CDK4*. These findings suggest genotype-specific mechanisms of RB pathway misregulation in GBM that would not have been apparent from genomic characterization of a tumor sample and could be differentially targeted during treatment. Our findings thus warrant further investigation to identify the mechanistic differences and non-redundant roles between CDK4 and CDK6 in GSCs to explain these genotype-specific dependencies. While our ability to uncover additional patient-specific vulnerabilities is currently limited by the relatively small number of genome-wide screens performed in patient-derived GSCs sharing the same genomic alterations, additional efforts will provide the required power. Given the heterogeneity of GBM, it is very unlikely that a single therapeutic strategy will be universally successful, and thus we foresee that patient stratification based on genotype-specific vulnerabilities will be an area of important focus for future studies, particularly if they are superimposed on common vulnerable pathways.

A comparison of our essentiality screens performed on freshly isolated GSCs from patients to screens conducted in serum-cultured glioma cell lines (Meyers et al., 2017), revealed striking differences in essentiality profiles, with the GSC screens described herein uniquely identifying genes involved in cell fate determination and gliogenesis. These findings highlight the strength of our GBM cell model which captures the stem cell characteristics of the disease. Consistent with this notion, we identified a substantial number of master neurodevelopmental transcription factor genes that have been linked to GBM stemness, including an interesting core gene network that was described as being capable of reprogramming differentiated GBM cells into a stem cell state (Suvà et al., 2014). *SOX2* and *SOX9*, in particular, were identified as essential genes across patient GSC cultures. SOX2 has been described to be a core GBM reprogramming factor and SOX9 has been shown to have an important role in maintaining neural stem cells, regulating astrogliogenesis (Caiazzo et al., 2015; Scott et al., 2010) and sphere formation in serum cultured glioma cell lines (Wang et al., 2018). Based on these studies and our screen results, SOX2 and SOX9 likely have foundational roles in driving GBM progression and maintaining stem cell properties in GSCs. Furthermore, we also identified genetic dependencies on additional SOX transcription factors that were variable across GSC cultures. Further studies are needed to help elucidate the mechanisms underlying how these neural stem cell genes are co-opted in GBM and how they work together to maintain the tumor initiating stem cell fraction. Although these transcription factors are notoriously difficult to target using small molecule approaches, these studies reinforce a need to continue to derive strategies that target the core GBM developmental/stemness state. Importantly, we also uncovered additional genes that may contribute to GBM stemness function, such as *SOCS3*. In contrast to a previous report of SOCS3’s predominant tumor suppressive function via negative regulation of JAK/STAT signaling (Croker et al., 2008), our data support a context-specific role as a positive regulator of GSC function, consistent with other reports showing SOCS3 enhances GBM tumor cell survival and treatment resistance (Zhou et al., 2007).

Recent advances in the field of epigenetics demonstrate the importance of epigenetic regulation in determining cell fate (Hsieh and Zhao, 2016; Podobinska et al., 2017). We identified several epigenetic modulators as unique essential genes in GSCs. We studied the histone methyltransferase DOT1L in depth due to its conserved essentiality. Recently, DOT1L was identified as an *in vivo*-specific genetic dependency in GBM using a focused RNAi screen (Miller *et al*., 2017). Intriguingly, shRNA-mediated knockdown of *DOT1L* was deemed dispensable for the growth of GSCs *in vitro,* while we found that CRISPR-mediated knockout of *DOT1L* robustly inhibits the growth of all GSC cultures. The reason for this discrepancy is unclear but may point to residual DOT1L activity due to incomplete knockdown obtained with shRNA or to the delay in observable effect upon DOT1L loss of function. Nevertheless, together with the previous report, the evidence strongly points to the importance of epigenetic control in regulation of functional stem cell properties. We extend the previous findings by providing evidence that two GSC essential genes, SOX2 and OLIG2, are epigenetically regulated by DOT1L histone methylation, suggesting epigenetic mediated regulation, in part is involved in targeting core stemness programs. Development of brain-permeable DOT1L inhibitors or drug delivery modalities to enable inhibitors such as EPZ-5676 to access the tumors will be worthwhile for pre-clinical testing. Overall, the data support the continued design of therapeutics that specifically exploit epigenetic vulnerabilities in order to access GBM stemness function.

One of the more exciting findings from our screens was the identification of multiple stress signaling pathways as essential for the growth of GSCs. The top GSC-specific hit in this study, c-JUN, plays an important role in the protection of cells from stress-induced apoptosis. Supporting these results, the kinase upstream to the JNK-JUN signaling axis, MAP2K7 was also identified as essential. Together with previous studies (Matsuda et al., 2012; Yoon et al., 2012), our findings support further investigation of the effectiveness of therapeutic strategies targeting JNK-JUN signaling for GBM. Along the same lines, the identification of several components of the protein ufmylation pathway as essential genes, further highlights the importance of stress response mechanisms for the survival and/or growth of GSCs. Indeed, given that this ubiquitin-like post-translational modification is linked to modulation of the ER stress response (Zhang et al., 2012), these results suggest that GSCs are under high proteotoxic stress and addicted to ER stress signaling pathways, rendering them vulnerable to perturbation of proteostasis gene networks (Deshaies, 2014). This finding is in line with new studies describing unique molecular mechanisms employed by stem cells or their malignant counterparts to maintain their unique functional properties (Dolma et al., 2016; van Galen et al., 2014; Tang and Rando, 2014). Our findings therefore provide a molecular basis for triggering proteotoxic crisis (Urra et al., 2017), either by inhibiting stress response pathways or by increasing ER stress over homeostatic levels, such as by inhibition of proteasome activity (Raizer et al., 2016), as a consideration for GBM treatment.

To complement our efforts to uncover new genetic vulnerabilities in GBM, we also performed chemogenomic screens to better understand the genes and pathways that govern the variable responsiveness of GSCs to TMZ. We also aimed to identify targets for combination therapy to increase the efficacy of chemotherapy for GBM patients by blocking mechanisms of intrinsic TMZ resistance. Positive selection screens in two GSC cultures identified just four genes, all core members of the MMR pathway, whose loss confers resistance to high dose TMZ, faithfully modeling acquired TMZ resistance observed in GBM recurrence (Wang et al., 2016). More importantly, from a therapeutic perspective, our negative selection screens with sub-lethal TMZ doses revealed numerous genes underlying intrinsic TMZ resistance whose loss of function mutations increase sensitivity to TMZ. Most of these genes converge on the Fanconi Anemia/interstrand crosslink repair and homologous recombination pathways. Supporting these results, several studies have linked the Fanconi Anemia pathway to TMZ resistance (Chen et al., 2007; Kondo et al., 2011) Our screens also identify multiple genes involved in Base Excision Repair (BER) and Nucleotide Excision Repair (NER), such as *APEX1* and *MPG*, consistent with TMZ-induced DNA adducts also being BER substrates. These hits were however not conserved across all GSC cultures indicating variable activity or engagement of resistance pathways dependent on patients. Nevertheless, many of the conserved hits have potentially druggable enzymatic functions including, for example, *MCM8* and *MCM9* which we validated to result in TMZ hypersensitivity when knocked out. In addition to genes with known functions in DNA damage repair, we also identified several uncharacterized factors, such as the zinc finger domain containing protein ZC3H7A, which we describe to play a role in GSC resistance to TMZ.

In summary, our genome-wide CRISPR screens in patient-derived GSC cultures have identified a diversity of genetic vulnerabilities in GBM tumorigenic cells. We identify a substantial number of pathways that previously were not linked to GBM biology, but now can be nominated for further preclinical exploration. As highlighted here, we suggest that advances in GBM therapies will need to combine in depth genomic characterization with functional approaches to unveil the genes and pathways functionally connected to properties of the key malignant population. Despite the concern for tremendous interpatient heterogeneity and the implications of nihilism that this view brings to future therapeutic development, the finding of common essential programs for GSC growth and survival lends hope that we will develop therapies that can be used on larger groups of GBM patients, with patient specific vulnerabilities providing complementary personalized strategies.

## Acknowledgments

Research was supported by SU2C Canada Cancer Stem Cell Dream Team Research Funding (SU2C-AACR-DT-19-15) provided by the Government of Canada through Genome Canada and the Canadian Institutes of Health Research, with supplementary support from the Ontario Institute for Cancer Research through funding provided by the Government of Ontario. Stand Up To Cancer Canada is a program of the Entertainment Industry Foundation Canada. Research funding is administered by the American Association for Cancer Research International – Canada, the scientific partner of SU2C Canada.

S.A. was also supported by the Canadian Institute of Health Research and the Terry Fox Research Institute.

P.B.D. is also supported by the Terry Fox Research Institute, the Canadian Cancer Society, the Hospital for Sick Children Foundation, Jessica’s Footprint Foundation, the Hopeful Minds Foundation, the Bresler family, and B.R.A.I.N. Child. P.B.D. holds a Garron Family Chair in Childhood Cancer Research at The Hospital for Sick Children.

S.W. and H.A.L. are also supported by the Canadian Institutes of Health Research.

T.H. was supported by MD Anderson Cancer Center Support Grant P30 CA016672 (the Bioinformatics Shared Resource) and the Cancer Prevention Research Institute of Texas (CPRIT) grant RR160032.

We thank Dr. Mark Bernstein (University Health Network, Toronto), and Dr. Sunit Das and Dr. Michael D Cusimano (St. Michael’s Hospital, Toronto) for the generous provision of tumor tissue. In addition, we thank Dr. Alice Yijun Wang for help in data acquisition and analysis, Dr. Michael Johnston for feedback on the manuscript, Rozina Hassam and Orsolya Cseh (University of Calgary) for technical support and Dr. Ahmed Aman (Ontario Institute for Cancer Research) for the mass spectrometry analysis for pharmacokinetic studies.

## Author Contributions

S.A, P.B.D. and G.M. conceptualized the study. D.A.B., I.R., X.H., H.A.L. and S.W. conceptualized the validation of DOT1L. G.M. and N.R. performed the genome-wide CRISPR-Cas9 screens. T.H., G.M., Z.S. and S.A. analyzed screen data. G.M., D.A.B., V.M., N.R., M.M.K., M.A., and H.Y. performed *in vitro* experiments. F.C. contributed to the drug response experiments in GSCs. D.A.B., X.H. and H.A.L. performed *in vivo* experiments. P.B.D., H.A.L. and S.W. provided the GBM culture. S.A., P.B.D., H.A.L. and S.W. supervised the study and acquired funding for research. S.A. P.B.D. and G.M. wrote the manuscript with input from all authors.

## STAR Methods

### Contact for Reagent and Resource Sharing

Additional data and requests for resources should be directed to the lead contact. Materials can be obtained via material transfer agreement from authors institutions upon reasonable request to corresponding authors.

### Experimental Model and Subject Details

#### Cell Culture

Glioblastoma stem cells (G432NS, G472NS, G477NS, G523NS, G510NS, G440NS, G564NS, G361NS and G567NS) and human neural stem cells were grown on poly-l-ornithine (Sigma-Aldrich) and laminin (Sigma-Aldrich) coated Primaria cell culture dishes (Corning) in Neurocult NS-A Basal media (StemCell Technologies) supplemented with N2 (20ng/mL, Life Technologies), B27 (Life Technologies), EGF (10ng/mL Life Technologies), FGF (10ng/mL Life Technologies), heparin (2ug/mL), BSA (150 µg/mL) and GlutaMAX (Life Technologies) as described by Pollard *et al*. (Pollard et al., 2009). Cells were dissociated using Accutase (Life Technologies). BT67, BT119, BT147, BT89 and BT69 GSCs were cultured in neurosphere conditions on non-adherent plates in serum free media supplemented with EGF (20 ng/mL; Peprotech, FGF2 (20 ng/mL; R&D Systems Inc) and heparin sulfate (2 µg/mL; R&D Systems). HEK293 T-REX Flp-in and Hela cells were grown in DMEM supplemented with 10% FBS. Where applicable, blasticidin, hygromycin and puromycin antibiotics were used at concentrations of 5 µg/mL, 200 µg/mL and 2 µg/mL respectively. GSC lines were confirmed to match their parental primary GBM tumor tissue by microsatellite genotyping (The Centre for Applied Genomics, Hospital for Sick Children) or short tandem repeat profiling (Calgary Laboratory Services and Department of Pathology and Laboratory Medicine, University of Calgary). Authentication and testing of all cell lines was performed as per American Association for Cancer Research recommendations.

#### Lentivirus production

Lentivirus containing the TKO gRNA library was prepared as previously described (Hart *et al.,* 2015) and concentrated using Lenti-X Concentrator solution (Clontech) according to the manufacturer’s protocol. For small-scale lentivirus production of individual gRNA constructs, approximately 3.5 million HEK293T cells were seeded on 10 cm plates 24 hours prior to transfection with 6 µg of lentiviral vector, 6 µg of psPAX2 and 2 µg of pMD2.G per plate using the calcium phosphate method. Viral media was harvested 48 and 72 hours post-transfection, cleared via centrifugation at 1,000 x g for 5 minutes and filtered prior to concentration using Lenti-X concentrator.

#### Generation of stable Cas9 expressing GBM cell lines

Patient-derived GSC lines G361NS, G432NS, G440NS, G472NS, G477NS, G510NS, G523NS, G564NS were infected with lenti-FLAG-Cas9-2A-BsdR virus (Hart *et al.,* 2015) in the presence of 0.8µg/mL polybrene and seeded onto PLO/laminin coated tissue culture dishes. Twenty-four hours post-transduction, media was changed and transduced cells selected with 2-5µg/mL blasticidin. CRISPR-Cas9 genome editing capability of GSC cells was tested using a validated lentivirus expressing pLCKO-gRNA-PSMD1 (5’-accagagccacaataagcca-3’) and compared to the negative control gRNA for LacZ (5’-CCCGAATCTCTATCGTGCGG-3’). PSMD1 Is essential for survival and thus cells that have undergone gene editing at the *PSMD1* locus die within a few days. GSC-Cas9 cells were transduced with lenti-pLCKO-gRNA-PSMD1 and selected with 2 µg/mL puromycin for 2 days followed by seeding at low density in 6 well plates and grown until the control well was confluent. Cells were then fixed in methanol and stained with 0.1% Crystal Violet. gRNA-PSMD1 wells displayed no visible viable cells indicating efficient gene editing.

#### Mouse Housing and Husbandry

6- to 8-week-old female CB-17 SCID mice (Charles River) were used. Mice were housed in a Biohazard barrier level 2 facility in “Techniplast Greenline” ventilated cages at a maximum density of 5 mice per cage for the duration of the study. Food and water was administered *ad libitum.* Animal care guidelines are as per the University of Calgary animal care guidelines. The University of Calgary is accredited by the Canadian Council on Animal Care.

#### Pharmacokinetic Analysis

SCID mice (N=6) were treated with 35 mg/kg by subcutaneous injection. Treatment was administered twice daily for 5 days. Control mice were treated with vehicle (5% hydroxypropyl-β-cyclodextrin (HPBCD) in saline and one mouse received no treatment. On the fifth day, 30 minutes after the first treatment of EPZ-5676, 3 mice from each treatment group were sacrificed and brains as well as plasma was collected. The untreated mouse was also sacrificed at this time point and brain and plasma collected. 5 hours following treatment, the remaining mice were sacrificed, and brains and plasma collected. Liquid-chromatography-mass spectrometry (LC-MS) was used to determine the concentration of drug accumulation in both the plasma and brain samples at both collection time points.

#### Flank Xenograft Studies

1×10^6^ dissociated BT147 GSCs were injected into the flanks of 16 SCID mice. Mice were then randomized into the vehicle or EPZ-5676 treatment groups. Mice were treated twice daily with vehicle (5% HPBCD in saline) or 35 mg/kg EPZ-5676. Once flank tumors reached a maximum diameter of 20 mm, mice were sacrificed, tumors were extracted and flash frozen with liquid nitrogen. RNA and protein were collected from 3 tumors from each treatment group to assess the effect of treatment on the expression of genes associated with neural and glial differentiation as well as DOT1L histone methylation, respectively.

#### Orthotopic Xenograft Studies

Mice were anesthetized intraperitoneally with a mixture of ketamine (200 mg/kg) and xylazine (10 mg/kg). A one cm incision was made on the skin above the skull. Using a Kofp stereotactic apparatus with a mouse adapter, a burr hole was then drilled into the skull of each mouse. Luciferase labelled (Campeau et al., 2009) (Addgene #21474) GSCs were pre-treated with DMSO or 1 µM EPZ-5676 for 28 days. Cells were dissociated, and drug replaced every 7 days. 50 000 viable BT147 cells and 100 000 viable BT67 cells (as assessed with Trypan Blue exclusion) were injected into the right cerebral hemisphere at a depth of 3.5mm using a 10 L hamilton syringe. The incision was closed with surgical staples. Post-surgery, animals were monitored during their recovery from anesthesia then returned to the normal housing area. Every 7 days mice were injected intraperitoneally with 150 mg/kg body weight luciferin (Gold Biotechnology) and tumor burden assessed with a Xenogen IVIS 200 Spectrum imager. tumor size was quantified with Living Image software from PerkinElmer. Mice were imaged until they exhibited significant weight loss or signs of neurological symptoms such as severe hunching, domed heads and/or lack of grooming. Mice were then sacrificed, and end-point recorded for each mouse. Survival data was plotted and analyzed using GraphPad Prism.

### Methods Details

#### Genome-wide CRISPR-Cas9 screens

CRISPR-Cas9 screens were performed using the 90K TKOv1 library (Hart et al., 2015) (Addgene #1000000069). A minimum of 60×10^7^ GSC-Cas9 cells (cultured with 2-5µg/mL blasticidin) were transduced at an M.O.I. of 0.3-0.4 with the TKO library for minimum 200-fold coverage, in the presence of 0.8µg/mL polybrene and seeded onto 30-40 10 cm dishes (seeding density was cell line dependent). Twenty-four hours post-transduction, media was changed and replaced with fresh GBM media containing 2-5µg/mL blasticidin and 2µg/mL puromycin. After 48 hours of puromycin selection, cells were combined and two 1.8×107 cell T0 samples were collected and stored at −80°C for later processing. Remaining cells (minimum 1.8×10^7^) were split into biological replicates and seeded across 10 cm dishes for a further 5 days to allow for efficient genome editing. At this point (7 days post-transduction) cells were divided into 2-4 replicates (depending on concurrent TMZ screens) each containing 1.8×10^7^ cell seeded across ten to fifteen 10 cm dishes. For TMZ hypersensitivity screens, two replicates were treated with an experimentally determined sub-lethal (LD10-20) dose of TMZ and two replicates treated with 0.02% DMSO. For TMZ resistance screens, two replicates were treated with an experimentally determined lethal dose (LD90) of TMZ. Replicate cell populations were cultured as such in the absence of antibiotics for a total of fourteen cell doublings of control replicates with fresh media/drug added every 2-3 days and passaging as needed (generally every 4-5 days, maintained at minimum of 200-fold library coverage). At the conclusion of the screen cell replicates were collected in 1.8×10^7^ cell aliquots (200-fold coverage).

#### Genomic DNA Library Preparation and Sequencing

Genomic DNA was extracted from frozen cell pellets using the QiaAMP DNA Blood Maxi Kit (Qiagen) using the manufacturer’s protocol, ethanol precipitated and resuspended at a concentration of 400 ng/µL. For each gDNA sample, 50µg of gDNA was amplified in ten 50µL PCR reactions using the KAPA HiFi Master Mix (Kapa Biosystems) and 1µM of “outer” pLCKO library primers (forward primer 5’-agggcctatttcccatgattcctt-3’; reverse primer 5’-tcaaaaaagcaccgactcgg-3’) for 19 amplification cycles. Individual sets of ten identical outer PCR reactions were pooled and 2.5µL was used as a template for a subsequent PCR reaction to add TrueSeq adaptor sequences and unique combination of i5 and i7 barcode sequences for each sample. For each sample 2 identical 50µL PCR reactions were performed using KAPA HiFi Master Mix and 1 µM of barcode primers for 17 amplification cycles. 200 bp barcoded PCR product was gel purified and submitted for next generation sequencing using Illumina HiSeq2500s or NextSeq500 instruments. T0, positive selection and negative selection samples were sequenced at read depths of 500-, 100- and 200-fold respectively.

#### CRISPR-Cas9 gene editing

For single gene knockout studies, individual gRNAs were ligated into *BfuAI* digested pLCKO, or *BsmBI* digested pLentiguide NLS-GFP/mCherry-2A-PURO (generous gift from Dr Daniel Durocher, Lunenfeld-Tanenbaum Research Institute, Toronto). GSC-Cas9 cells were infected with pLCKO-gRNA/pLentiguide-NLS-GFP-gRNA-2A-PURO lentivirus as above and selected with puromycin for 2-3 days prior to seeding in viability/drug treatment assays or MCAs. Gene editing was quantitatively assessed using TIDE (Brinkman et al., 2014) from PCR amplicons flanking gRNA target sites. For a complete list of gRNA oligo sequences, see Table S7.

#### Multicolour Competition Assay

To validate gene knockouts producing fitness defects multicolour competition assays (MCA) were conducted wherein equivalent number of GSC-Cas9 cells co-expressing NLS-mCherry and a non-targeting control gRNA were co-cultured with cells expressing NLS-eGFP and a gRNA targeting the gene of interest, and then were allowed to grow and compete during a 3-week period. GSC-Cas9 and hNS-Cas9 cells were transduced with lentivirus expressing either Lentiguide-LacZ-NLS-mCherry-2A-PURO or Lentiguide-gRNA-NLS-GFP-2A-PURO containing a gRNA to a target gene of interest and selected for integration with puromycin. To start an assay gRNA-LacZ-mCherry cells were mixed 1:1 with gRNA-geneX-GFP cells in a 12-well plate (10,000 cells per construct). One day after seeding (T0) wells were imaged with an InCell6000 or Cytation5 (BioTek) with a 4X objective and number of mCherry and GFP positive nuclei were counted from a montage of images covering most of the well using the Gen5 software (BioTek). Relative cell fitness was calculated from the percentage of GFP positive cells normalized to the T0 value.

#### Cell viability assays

GSC/GSC-Cas9/hNS cells were seeded in 96 well plates at a density of 1,000-3,000 cells per well (cell line dependent) in GSC media. Twenty-four hours following seeding, cells were treated with the indicated drugs/compounds. TMZ dose response experiments in gene edited GSC-Cas9 cells were conducted using a range of 0-100 µM of TMZ for 7 days with one media change with new drug added at mid-point. DKM 2-93 (MedChemexpress), SP600125 (Sigma-Aldrich) and Bentamapimod (Cayman Chemical) treatments were performed at indicated concentrations for 7 days. For MCM8 rescue experiment, pSNT-AAVS1-MCM8 was generated by replacing AscI/PacI fragment of pSNT-AAVS1-Cas9-2A-BFP (a generous gift from Dr. Ajamete Kaykas, Novartis) with full-length MCM8 human cDNA sequence, producing a Dox-inducible MCM8 expression construct. pSNT-AAVS1-MCM8 was transfected into G361NS-Cas9 cells along with a gRNA targeting the AAVS1 safe-harbour site and MCM8 integrated clones were selected with G418. MCM8 expression was induced with 1 µg/mL Dox for 48 hours prior to TMZ treatment and maintained throughout cell viability experiments with fresh Dox every 3 days.

For viability assays in sphere grown cultures (G523NS, BT147, BT119, BT89, BT67, BT69), GSCs were dissociated with Accumax (eBioscience) and 1000 cells in 100 µl growth factor containing media were plated in 96-well plates. Cells were left to recover for 24 hours before drug re-suspended in dimethyl sulfoxide (DMSO) was added in another 100 µl of media. Cells were treated with EPZ-5676 (Epizyme) at concentrations ranging from 0.01 µM to 10 µM with 6 wells per drug concentration. Control treated cells were left untreated or treated with the same volume of DMSO as the highest concentration of drug treated cells. At experiment endpoints, cell viability was assessed using the alamarBlue^TM^ reagent (Thermofisher) according to the manufacturer’s protocol. Results represent means from 3 biological replicates, each containing two technical replicates. Statistical analysis conducted using a two-tailed student’s t-test. DKM 2-93 experiments in GSC screening panel are mean of 2 biological replicates, each containing two technical replicates.

#### Immunofluorescence Microscopy

Cells were fixed on glass coverslips in 4% Paraformaldehyde. Where appropriate, cell permeabilization in 0.25% Triton-X PBS was performed for 10 minutes at room temperature followed by blocking in 0.1% Triton-X plus 1% Goat Serum in PBS for 60 minutes at room temperature. Primary antibody incubations were performed overnight at 4°C, followed by extensive washing. Primary antibodies used in this study include: TUBB3 (TU-20, Millipore-Sigma), Ki67 (SP6, Thermofisher) and γH2AX (JBW301, Millipore-Sigma). Secondary antibody incubations were performed for 1-2 hours at room temperature in the dark. Cells were mounted in Vectashield (Vector Labs) plus DAPI. Images were captured using either a Plan-Apochromat 20x/0.8 numerical aperture or 63×/1.4 numerical aperture oil immersion objective on a confocal microscope (LSM 700, Carl Zeiss) operated with ZEN software black edition. EGFP/Alexa Fluor 488 and mCherry fluorophores were excited individually with 488 and 543-nm lasers, respectively, with appropriate filter sets.

#### TUBB3 and Ki67 quantitation

Following six days of puromycin selection to allow time for efficient gene editing, pLentiguide-NLS-GFP-gRNA-*LacZ*-2A-PURO or pLentiguide-NLS-GFP-gRNA-*SOCS3*-2A-PURO transduced G510NS-Cas9 cells were fixed and processed for immunofluorescence microscopy as indicated above. Raw images were adjusted for brightness and contrast. Total cell number was quantified using the ImageJ software. TUBB3 positive cells were scored based on detection of fluorescent signal throughout the cytosol. Ki67 positive cells were scored based on nuclear staining. Statistical analysis was performed using the GraphPad Prism 5 software and significance was calculated using a paired, two-sided t-test from 3 biological replicate experiments.

#### Sphere Assays

Sphere assays were used to assess the effect of EPZ-5676 on GSC sphere formation and morphology. Cells were treated the same way as specified above for viability assays and following 14 days of treatment, 96-well plates were photographed using an Incucyte Live Imaging System (Essen Bioscience). To verify findings, 3 biological replicates were performed for each cell line.

#### Histone Extractions

Histone extractions were performed using a protocol that was adapted from Lu *et al.* and Schehter *et al* (Lu et al., 2012) (Shechter et al., 2007). Treated GSCs were pelleted and flash frozen with liquid nitrogen and stored at −80°C. Pellets were then lysed in hypotonic lysis buffer (10 mM HEPES, 10 mM KCl, 1.5 mM MgCl_2_, protease inhibitors (Roche) and 0.5 mM DTT) for 1 hour on ice and vortexed every 10 minutes. The cytosolic fraction was then removed, pellets were resuspended in cold 0.2N H2SO4 and rotated for 2 hours at 4°C. Nuclear debris was removed by centrifugation. Supernatant containing histones were transferred to new 1.5 mL Eppendorf tubes for trichloroacetic acid (TCA) precipitation, 33% TCA was added dropwise to the histone solution and incubated for 45-60 minutes on ice. Histones were then pelleted by centrifugation and washed twice with acetone. Histones were re-suspended in 60 µL distilled H2O with 4X Laemmli buffer and 10 µL 1 M Tris (pH 8).

#### Western Blotting

Primary antibodies were used to stain western blots overnight. Antibodies that were used include: H3K79me2 (3594, Abcam), H3K27me3 (6002, Abcam), H3K9me2 (1220, Abcam) H3 (10799, Abcam), TUBB3 (Millipore-Sigma) and GAPDH (Santa Cruz Biotechnology). Following incubation with primary antibodies, washed blots were incubated for an hour with horseradish peroxidase (HRP)-conjugated secondary antibodies. The secondary antibodies that were used include: donkey anti-mouse (Millipore) and goat anti-rabbit (Cell Signaling Technology). Blots were washed with tris-buffered saline and tween 20 (TTBS) (50 mM Tris, 150 mM NaCl, 0.05% Tween) and blocked with tris buffered saline (TBS). Blots were then imaged with SuperSignal West Pico chemiluminescent solution (Thermofisher) and Amersham Imager 600 (General Electric).

#### Quantitative Polymerase Chain Reaction (qPCR)

Treated GSCs were pelleted, flash frozen with liquid nitrogen and stored at-80°C. RNA was extracted using the RNeasy kit (Qiagen) following manufacturer’s instructions. Genomic DNA elimination was performed using Qiagen gDNA eliminator columns. RNA was then reverse transcribed using the qScript protocol (Quanta Biosciences), and cDNA was used for qPCR using the Lightcycler 96 standard protocol. See Table S8 for sequences of primers used to amplify *SOX2, OLIG2, NG2, TUBB3, NEUROD1, GFAP and ACTIN*.

#### Limiting Dilution assays

GSCs were pre-treated for 7 or 21 days with DMSO vehicle or 1 µM EPZ-5676. Cells were dissociated, and drug was replaced every 7 days. Cells were then dissociated and plated in 100 µL of growth factor containing media in a 96-well plate at densities ranging from 512 cells/well to 1 cell/well with 6 wells per cell density. Vehicle or drug was added to limiting dilution plates. Cells were left to form spheres for 14 days. The number of wells that exhibited sphere formation was then recorded for each cell density and results were analyzed using the Extreme Limiting Dilution Analysis computational program (Hu and Smyth, 2009). Average sphere initiating frequency as well as upper and lower 95% confidence intervals were plotted using GraphPad Prism. For each culture, 3 biological replicates were performed to verify results were consistent and significant.

#### Invasion Assays

Small GSC spheres were collected from culture flasks and treated in 6-well plates. Spheres were treated with DMSO or EPZ-5676 for 7 days. Spheres were then transferred to 1.5 mL Eppendorf tubes to allow gravity pellets to form. Media and treatment was removed, and spheres were resuspended in a Rat Collagen I (Cultrex) matrix supplemented with 10% FBS. 100 µL/well of collagen suspended spheres was then transferred to a cold 96-well plate with 3 experimental replicates for each treatment condition. Plates were transferred to an Incucyte Live Imaging System (Essen Bioscience). Area of cellular invasion was recorded hourly for several days before data was plotted using Graphpad Prism. Two-way ANOVA was used to determine statistical significance between treatment conditions for each GSC culture.

#### Migration Assays

GSCs were treated with DMSO or 1 µM EPZ-5676 for 7 days. Cells were then dissociated, and 2 500 cells resuspended in 50 µL non-growth factor supplemented media. Cells were plated in the top insert of a chemotaxis migration plate (Essen Bioscience) that was coated with a thin layer of Rat Collagen I matrix (Cultrex). 3 experimental replicates were plated per condition. In the bottom insert of the plate, growth factor supplemented media with 10% FBS was plated. The total area of migrated cells was measured hourly for several days using an Incucyte Live Imaging System (Essen Bioscience). Two-way ANOVA was used to determine statistical significance between treatment conditions for each GSC culture.

#### Flow Cytometry

GSCs were treated for 7 days with DMSO or 1 µM EPZ-5676. Cells were then dissociated and stained using anti-hNG2-APC (LHM-2, R&D Systems 130-105-329) and anti-GFAP-PE (REA335, Miltenyi Biotec FAB2585A) re-suspended in MACS/BSA (Miltenyi Biotec). To permeabilize, cells were resuspended in 2% ice cold MeOH and antibody was diluted in PBS (0.3% Triton). As controls for gating, Mouse IgG1-APC and REA Control (I)-PE human (Miltenyi Biotec) were used for NG2 and GFAP antibodies, respectively. Analysis and gating was done using FloJo software.

#### Proliferation and Cell Cycle Analysis

GSCs were treated with vehicle or EPZ-5676 for 7 and 21 days. Cells were dissociated, and drug was replaced every 7 days. Proliferation and cell cycle analysis was measured using the BrdU Flow kit from FITC BrdU Flow kit from BD Pharmingen (51-2354AK) following manufacturer’s instructions. BrdU was added to cells for 24 hours.

#### Apoptosis and Cell Death

GSCs were treated with vehicle or EPZ-5676 for 7 and 21 days. Cells were dissociated, and drug was replaced every 7 days. Apoptosis was measured using the Annexin V 568 kit (A13202) from BD Pharmingen following manufacturer’s instructions. Cell death was measured using LIVE/DEAD 488 fluorescent dye from Invitrogen (L34969).

#### H3K79me2 ChIP-qPCR

2.5 million dissociated GSCs were treated with DMSO or 1 µM EPZ-5676 for 5 days. Cells were then cross-linked with 1% formaldehyde for 10 minutes and glycine was added to a final concentration of 0.125 M to stop cross-linking. Cells were incubated on ice for 30 minutes in lysis buffer (1% SDS, 10 mM EDTA, 50 mM Tris-HCl and protease inhibitors (Roche) and sonicated for 45 minutes with a Biorupter to produce DNA fragments of 200-400 kb. Chromatin was then collected by centrifugation and flash frozen with liquid nitrogen for storage at −80°C. Magnetic beads (Dynabeads Protein A Life Technologies) were pre-washed with PBS BSA (5 mg/mL) and incubated overnight with primary antibodies H3K79me2 (3594, Abcam) or IgG (Cell Signaling Technology) at 4°C with rotation. Chromatin samples were divided for each antibody, 5 µL was used to verify sonication by running on an agarose gel and 5% of chromatin was saved for input. Samples were diluted 1:5 in dilution buffer (1% triton, 2 mM EDTA, 150 mM NaCl, 20 mM Tris-HCl and protease inhibitors (Roche) and incubated with magnetic beads pre-bound to primary antibodies overnight at 4°C with rotation. Magnetic beads were then washed with RIPA buffer (50 mM HEPES, 1 mM EDTA, 0.7% Na Deoxycholate, 1% NP-40 and 0.5 M LiCl) and 1 X TE buffer. Beads were re-suspended in elution buffer (1% SDS and 1% NaHCO3) and incubated at 65°C with vortexing every 30 minutes. Beads were then removed and immunoprecipitated chromatin as well as input samples were reverse cross-linked in elution buffer for 5 hours at 65°C with shaking. DNA was extracted using the QIAquick spin kit PCR purification kit following manufacturer’s instructions. DNA was used for qPCR using primers designed for the negative control heterochromatic regions QML5/6 as well as SOX2 and OLIG2. (Table S8). Three biological replicates were performed, and an unpaired t-test was used to determine significance for the changes in H3K79me2 levels between treatment conditions.

#### Co-localization studies

Hela or G361NS cells transduced with Venus-ZC3H7A lentivirus were transfected with mCherry-DCP1A or G3BP1 plasmids. mCherry-tagged G3BP1 and DCP1A (Youn et al., 2018) constructs were generous gifts from Dr. Anne-Claude Gingras (Lunenfeld-Tanenbaum Research Institute). 48 hours post-transfection cells were treated with 500 µM sodium arsenite or DMSO for 30 minutes at 37 °C and fixed in 4% paraformaldehyde for imaging as outlined above.

#### BioID Affinity Purification Mass Spectrometry

Full-length *ZC3H7A* cDNA was cloned into a pcDNA5/FRT/TO vector containing N-terminal hBirAm tag and used to create stable Flp-in T-Rex HEK293 cell lines as previously described (Coyaud et al., 2015). Expression of hBirAm-tagged ZC3H7A was induced in 2 x 15-cm dishes with 1 µg/mL tetracycline for 16 hours and followed by treatment with 25 µg/mL Biotin for 24 hours at which point cells were harvested and snap-frozen. Cells were thawed and lysed in RIPA buffer and insoluble material pelleted via centrifugation at 20,000 x g at 4° C. Cleared lysate was incubated with Streptavidin-Sepharose (GE Healthcare) overnight at 4°C with end over end rotation. Streptavidin resin was washed extensively with RIPA buffer and 50 mM ammonium bicarbonate and digested on-bead with sequencing grade trypsin. Digested peptides were resuspended in 0.1% formic acid, 5% acetonitrile in water. LC-MS/MS was performed on two biological replicates using an LTQ XL Linear Ion Trap Mass spectrometer (Thermo Scientific).

### Quantification and Statistical Analysis

#### Analysis of genome-wide screen data

Read counts for individual gRNAs from duplicate samples at each timepoint were mapped to gRNA library using MAGECK count function(Li et al., 2014). The BAGEL pipeline was used to filter out genes with <30 reads in T0 sample, normalize gRNA reads for sample sequencing depth, and calculate log2 fold-change relative to the T0 sample for the relevant screen. The BAGEL algorithm (Hart and Moffat, 2016) was then used to calculate Bayes Factor (BF) values for each gene, representing a confidence measure that the gene knockout results in a fitness defect. The algorithm uses reference sets of essential (n = 684) and nonessential (n = 927) genes (Hart et al., 2015). BAGEL software is available at https://github.com/hart-lab/bagel. To identify GBM-specific essential genes, raw BF scores from GBM and previously published TKO screens (Hart et al., 2015, 2017), (Zimmerman *et al* submitted) were quantile normalized and average normalized BF scores for GBM and non-GBM screens were calculated. Difference in average BF between GBM and non-GBM screens for each gene were transformed to Z-scores (ZBF) and Z-score ⪰2 was used as a cut-off score. Note that a previous analysis of G361NS essentiality screen, but not chemogenomics screens using a different set of reference essential genes was published in (Hart et al., 2015). For identification of GBM/Glioma specific essential genes in Toledo *et al.(Toledo et al., 2015)* GECKO and Meyers *et al. (Meyers et al., 2017)* Avana CRISPR-Cas9 screens were analyzed similarly with the exception of using Z-transformed CERES scores in place of BFs for Avana library screens. Complete linkage clustering of ZBF scores was performed using Heatmapper (Babicki et al., 2016) web server with the Euclidean distance measurement option. As a quality control measure, we included a F1-score cut-off of >0.75 at BF = 5 for screens to be included in the calculation of glioma/GBM specific essential genes. To quantify changes in BF between DMSO and TMZ treated samples in hypersensitivity screens, BF_control was subtracted from BF_TMZ for each gene in each individual screen and Z-scores were calculated independently for each screen.

### Gene Ontology Enrichment and Visualization

To determine Gene Ontology Biological Processes and Reactome Terms enriched in GBM-specific essential genes, all genes with ZBF ≥ 2 were used as input in gProfiler(Reimand et al., 2016) with recommended settings and filters (http://baderlab.org/Software/EnrichmentMap/GProfilerTutorial). Correction for multiple comparisons performed using Benjamini-Hochberg method. Resultant data was imported into Cytoscape using the Enrichment Map (Merico et al., 2011) app for visualization. Analysis of ZC3H7A BioID identified proteins was performed in the same manner with the addition of Gene Ontology Cellular Component terms. For Gene Ontology biological process and cellular component enrichment analysis in TMZ resistance screens, genes were ranked by TMZ sensitization Z-score and full ranked list was input into the GOrilla web application (Eden et al., 2009) and analyzed using default parameters.

#### Analysis of gene expression data

*SOCS3* expression levels and associated survival data was mined from TCGA and Rembrandt datasets using the GlioVis web interface (Bowman et al., 2017). Expression data depicted from Agilent-4502A platform TCGA dataset. For survival analysis, *SOCS3* high and low groups represent the highest 25% and lowest 25% expressing groups respectively with no sample exclusions based on CIMP status, *MGMT* status or recurrence. Statistical testing was performed using Gliovis interface via the log-rank method.

#### MS data acquisition and analysis

RAW files generated from LC-MS/MS analysis of BioID experiment were converted to .mgf format and uploaded to ProHits (Liu et al., 2010). Files were analyzed and searched using Mascot against the Human RefSeq V45 database consisting of forward and reverse “decoy” sequences. The Mascot version 2.3.02 (Matrix Science) search engine was used with trypsin specificity (two missed cleavages were allowed), deamidation (NQ), and oxidation as variable modifications. Additional parameters included charges of +2, and +3 allowed, precursor mass tolerance of 50 ppm, and fragment bin tolerance was set at 0.6 amu. Identified proteins having a minimum Mascot score of 50 and 2 unique peptides were filtered for common contaminants such as cytoskeletal proteins, keratins, artifact proteins, ribonucleoproteins and histones as well as proteins identified from the control set. SAINT analysis (Choi et al., 2010; Liu et al., 2010) using SAINTexpress was used to finalize the list of ZC3H7A interacting proteins with only proteins with a SAINT score of 1.0 considered.

### γH2AX-foci quantitation

For DNA Repair experiments, G361NS-Cas9 cells were seeded on PLO/Laminin coated coverslips and 16-24 hours later treated with 7.5 µM TMZ for 24 hours. At this point the media was replaced with fresh GSC media and cells were grown for 3 more days in the absence of TMZ followed by fixation with 4% paraformaldehyde and imaging as outlined above. Images were exported by ZEN software and analyzed using ImageJ to count γH2AX foci using the “Find Maxima” function with consistent noise level settings for all images. A minimum of 100 cells per replicate per condition were quantified for 2 biological replicate samples.

### Statistical Analysis

Statistical details of experiments can be found in figure legends. Unless otherwise indicated all statistical analyses were performed with Graphpad Prism 5 (GraphPad Software, La Jolla, California, USA). Unless otherwise indicated all statistical tests performed were two-sided. Statistical analysis of flow cytometry, BrdU, cell cycle and Annexin V data was performed using unpaired, two-sample t-tests from n=3 replicates. Statistical analysis of qPCR experiments was performed with one-way ANOVA from 3 biological replicates. For orthotopic xenograft experiments, survival was measured according to the Kaplan-Meier method, with the Mantel-Cox test used to compute statistical significance for 9 mice per treatment group for BT147 and 6 mice in the DMSO group and 5 in the EPZ-5676 group for BT67 (one mouse was excluded from the treatment as it had to be sacrificed early due to an eye infection). Statistical analysis of immunofluorescence microscopy images was performed in GraphPad Prism 5 (GraphPad Software, La Jolla, California, USA) and significance was calculated using unpaired, two-sample t-tests from n = 3 biological replicates. Statistical significance of cell viability and multicolour competition assays was assessed using an unpaired two-sample t-test from 3 biological replicates. Further details of individual experiments can be found in methods details.

---

**Figure S1.**
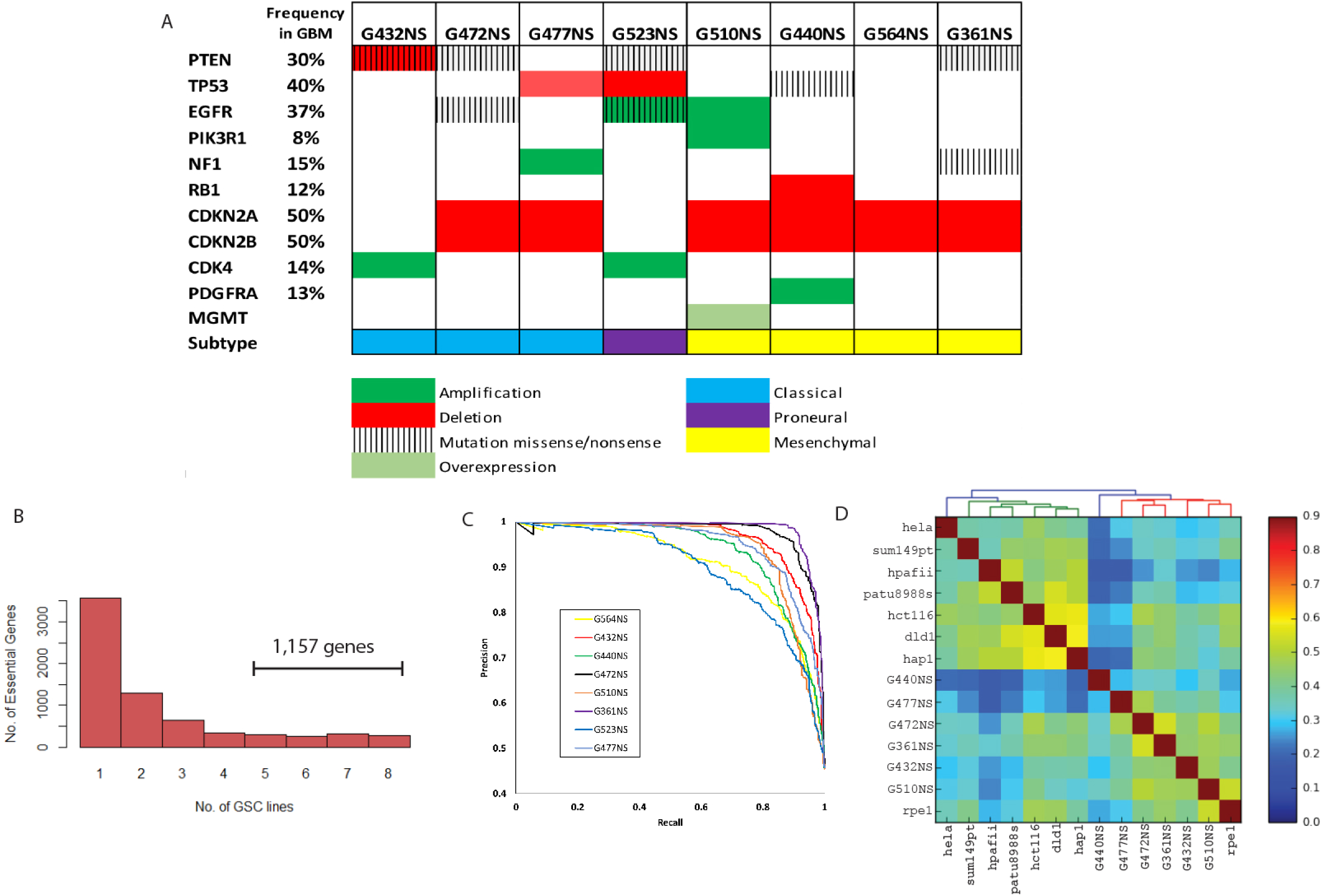
CRISPR-Cas9 screens supporting information. Related to Figure 1. (A) Genomic alterations in GSC CRISPR-Cas9 screening panel extracted from exome-seq,RNA-seq and CNV data. (B) Bar-plot showing the number of essential genes identified in 1-8 GSCs CRISPR-Cas9 screens. (C) Precision-recall curves for 8 GSC CRISPR-Cas9 screens produced with BAGEL pipeline and V2 reference essential/non-essential genes. Note sub-optimal performance of G523NS and G564NS screens. (D) Clustergram of Pearson correlation coefficients among all pairs of QC filtered TKOv1 screens.

**Figure S2.**
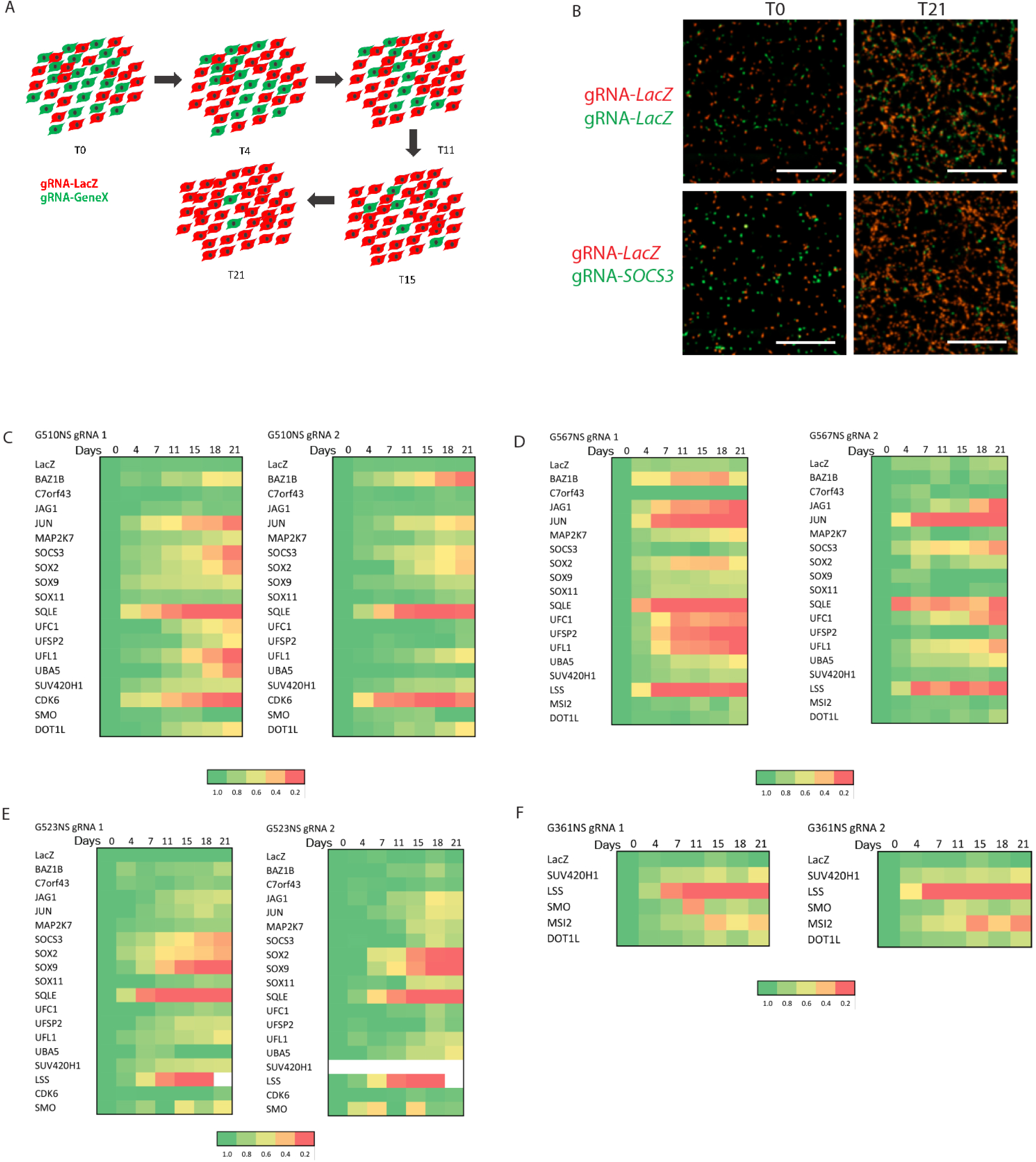
Validation of GBM essential genes. Related to Figure 1. (A) Schematic diagram for multicolour competition assay used to validate GBM essential genes. (B) Images from a Multicolour Competition Assay (MCA) in G510NS-Cas9 showing cells co-expressing a control gRNAs targeting *LacZ* and mCherry (Red) and cells co-expressing a gRNA targeting a candidate essential gene and GFP (green). Scale = 1 mm.(C-F) Heatmaps of MCA data for GBM essential genes across 4 GSC lines. Results represent averages from a minimum of n=3 (G510, G523) or n=2 (G567,G361) biological replicates per gRNA.

**Figure S3.**
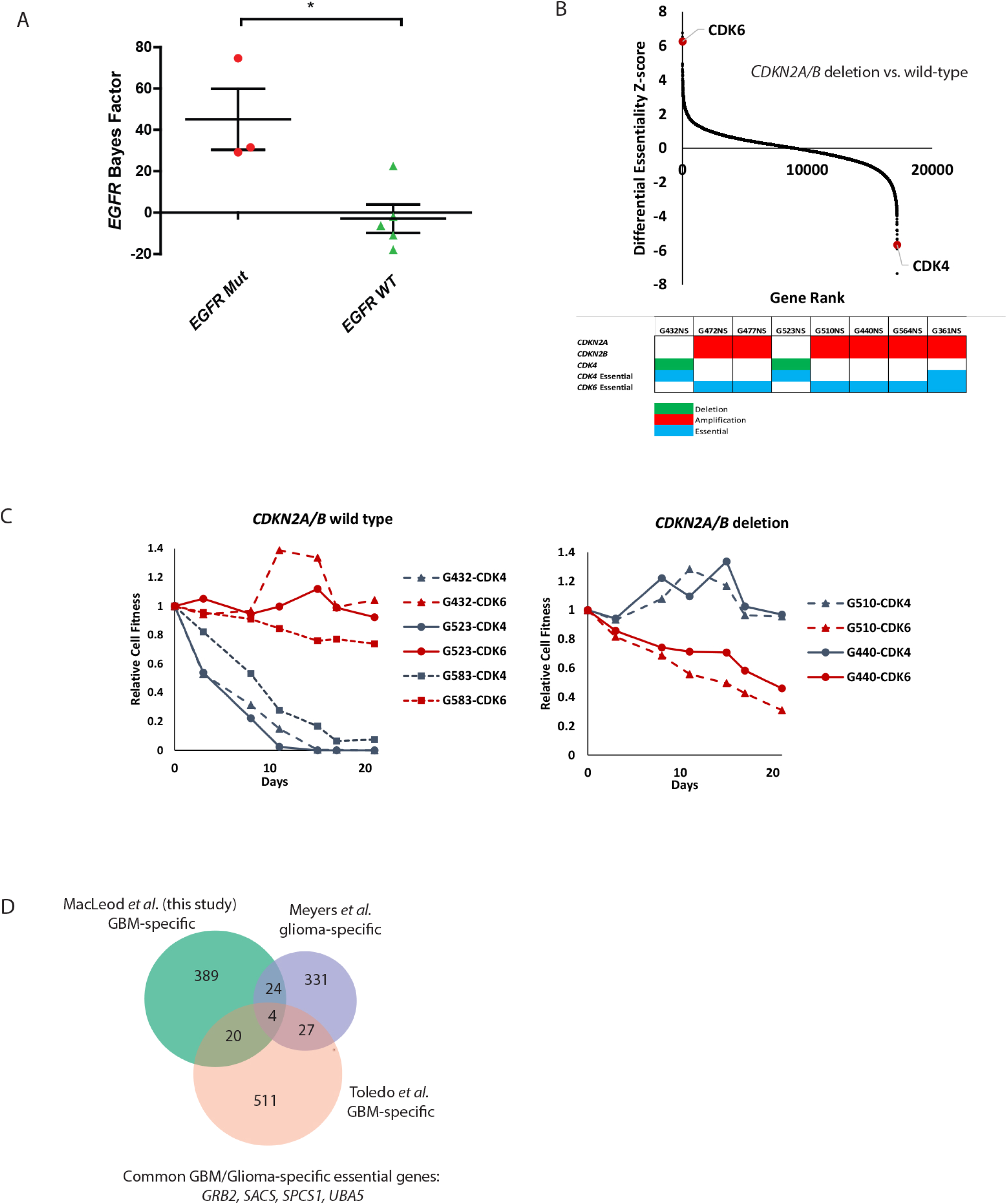
Context-specific essential genes in GBM. Related to Figure 1. (A) Bayes Factor scores for *EGFR* across 8 GBM CRISPR-Cas9 screens separated by mutational status into *EGFR* Mut (missense/nonsense mutation and/or gene amplification) or *EGFR* WT. Significance was assessed via a two-tailed t-test, * indicates p-value < 0.05. (B) Rank order plot comparing differential essentiality Z-scores in GBM cultures with *CDKN2A/B* deletion vs. WT highlighting the differential dependencies on *CDK6* and *CDK4* (top) with accompanying oncoprint/essentiality heatmap (bottom). (C) Multicolour competition assays using gRNAs targeting *CDK4* and *CDK6* in *CDKN2A/B* wild-type and deleted GBM cultures. (D) Overlap between GBM-specific essential genes (see Figure 1c), Glioma-specific essential genes from Meyers *et al.* 2017 and GBM-specific essential genes from Toledo *et al.* 2015.

**Figure S4:**
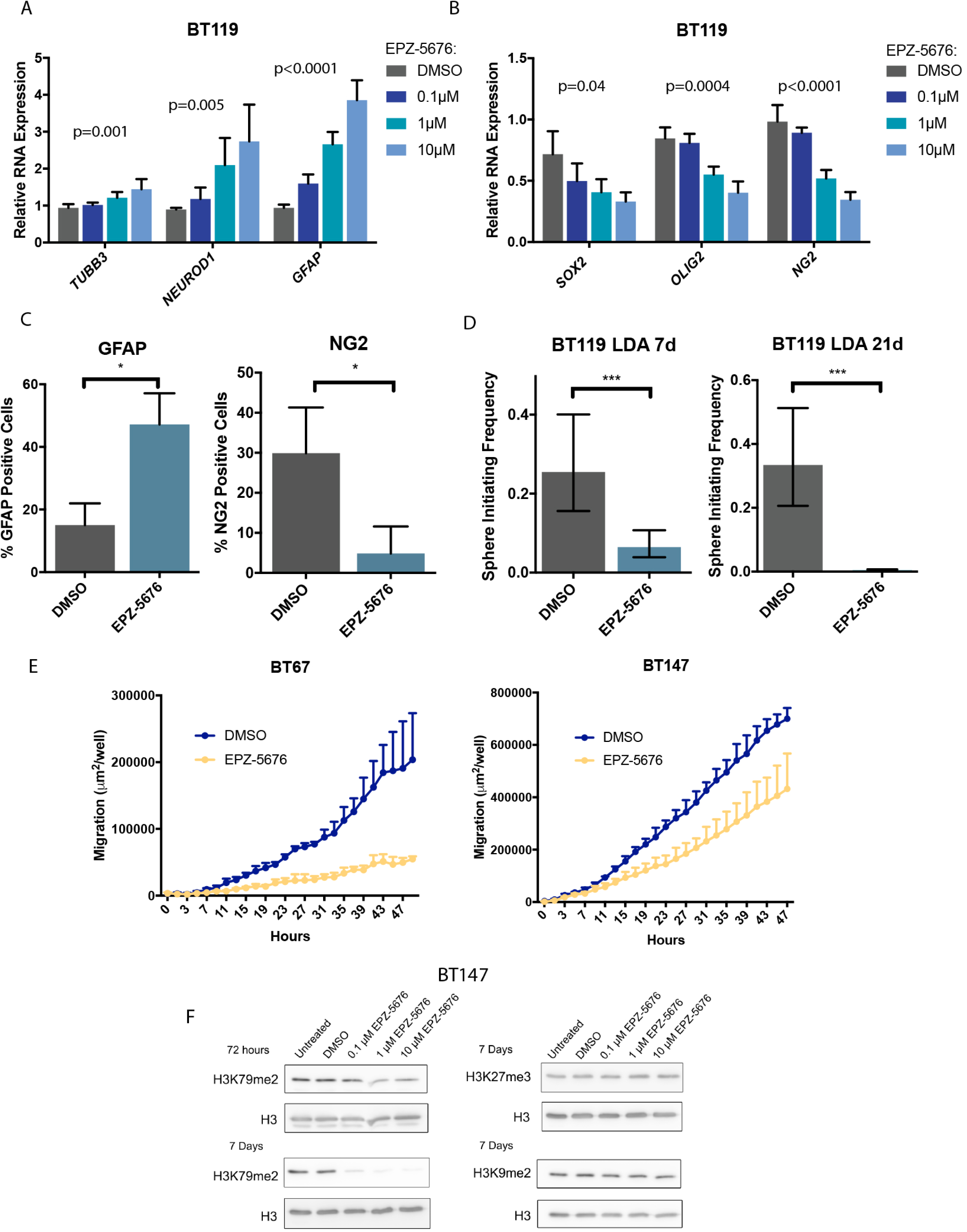
Validation of epigenetic regulation of DOT1L in GSCs *in vitro.* Related to Figure 3. (A) Gene expression of neuronal differentiation markers (*TUBB3* and *NEUROD1*) and astrocytic differentiation marker (*GFAP*) in BT119 GSCs treated with EPZ-5676. Data are represented as mean ± S.D. (B) Gene expression of stem cell marker (*SOX2*) and progenitor cell markers *(OLIG2* and *NG2*) in BT119 GSCs treated with EPZ-5676. Data are represented as mean ± S.D. (C) Protein expression of NG2 and GFAP as determined using flow cytometry in BT119 GSCs (unpaired t-test * p < 0.05). Data are represented as mean ± S.D. (D) Sphere initiating frequency of BT119 GSCs following short-term and long-term EPZ-5676 treatment. Data represents mean ± upper and lower 95% confidence intervals. (E) Quantified migration of GSCs following EPZ-5676 pre-treatment (Two-way ANOVA p-value BT67=0.005, p-value BT147=0.01). Data are represented as mean ± S.D. (F) Expression of H3K79me2 as well as H3K27me3 and H3K9me2 controls following EPZ-5676 treatment.

**Figure S5:**
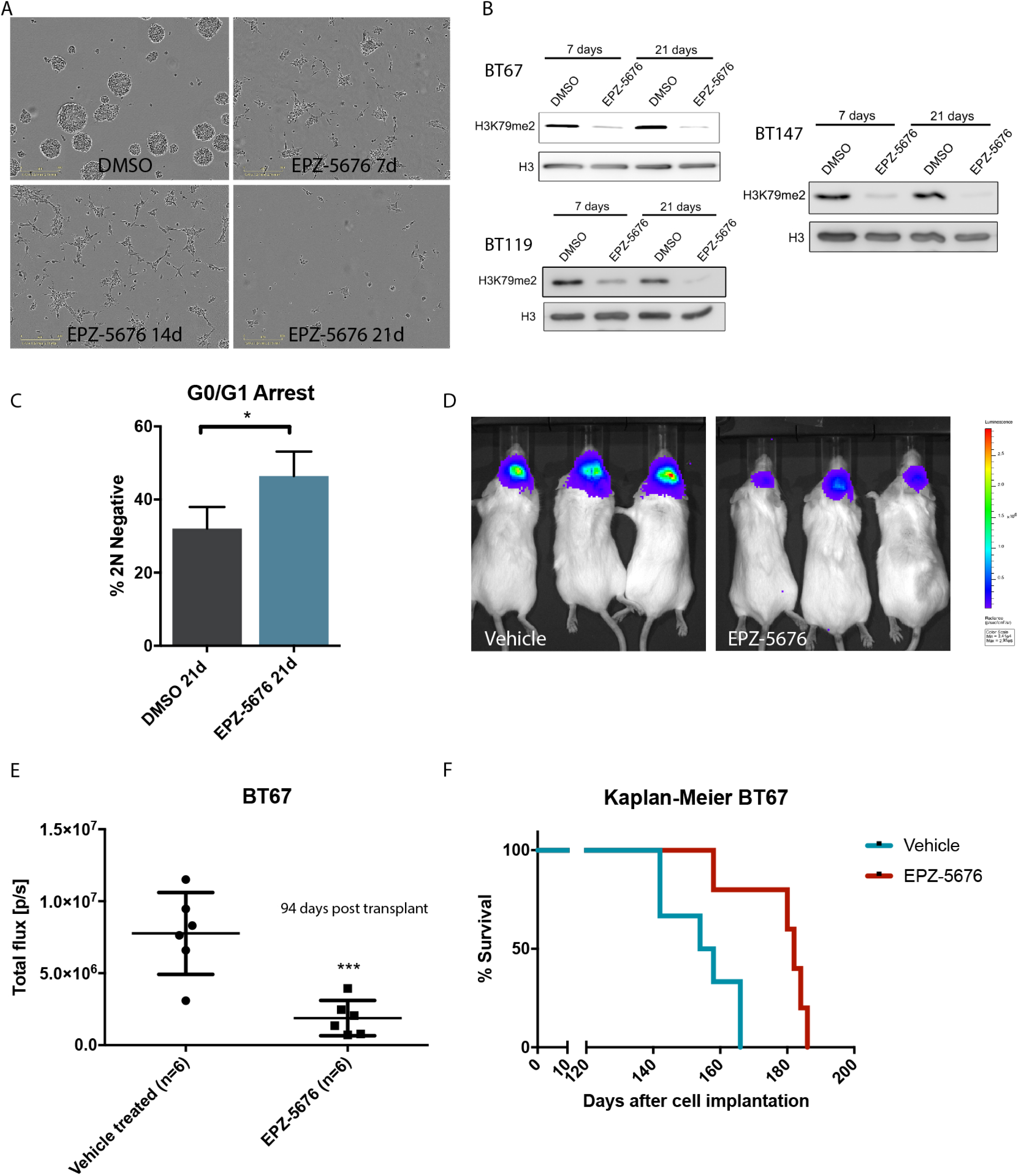
Effect of long-term DOT1L inhibition in GSCs *in vitro* and *in vivo.* Related to Figures 3 and 4. (A) BT67 GSC sphere formation following increasing treatment periods with EPZ-5676. (B) Expression of DOT1L-dependent H3K79me2 levels in 3 different GSC cultures following short-term and long-term treatment with EPZ-5676. (C) The effect of EPZ-5676 treatment on cell cycle (unpaired t-test * p < 0.05). Data are represented as mean ± S.D. (D) Representative tumor burden at 94 days post-engraftment for mice that were orthotopically xenografted with luciferase labelled BT67 that were pre-treated with vehicle or EPZ-5676. (E) Quantified tumor burden for mice that were orthotopically xenografted with BT67 that were pre-treated with vehicle or EPZ-5676 (unpaired t-test *** p < 0.001). (F) Kaplan-Meier survival curves for EPZ-5676 pre-treated BT67 in an orthotopic model (Mantel-Cox test * p < 0.05).

**Figure S6:**
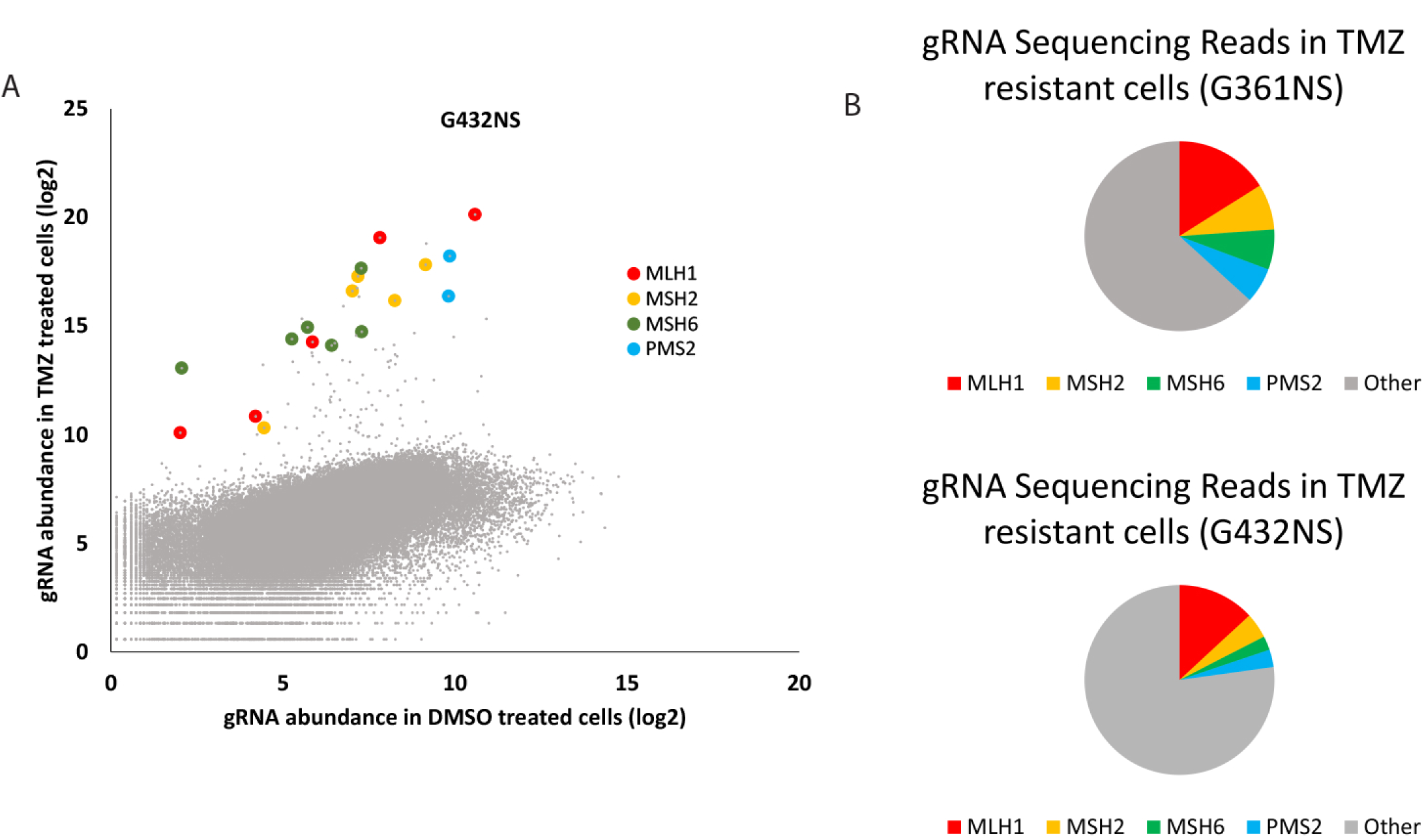
CRISPR-Cas9 positive selection TMZ screening additional data. Related to Figure 6. (A) Scatterplot of sequencing reads for gRNAs in G432NS TMZ sensitivity screen. Highlighted are gRNAs targeting(MMR genes *MLH1, MSH2, MSH6* and *PMS2*. (B) Representation of gRNAs mapping to MMR genes amongst total sequencing reads for endpoint samples in LD90 dose TMZ treated G361NS and G432NS lines in genome-wide screens.

**Figure S7:**
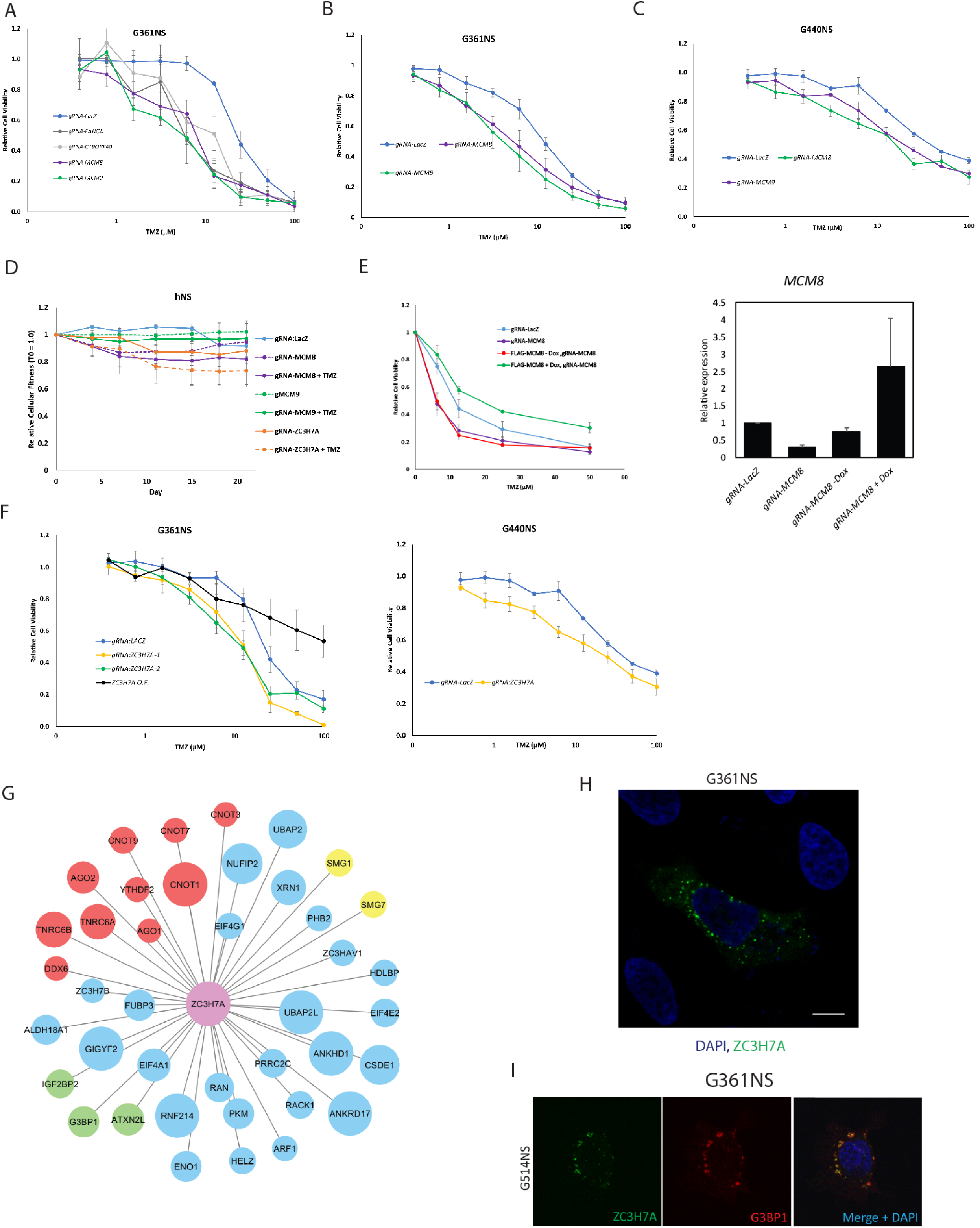
TMZ hypersensitivity knockouts extended data. Related to Figures 7. (A-C) Cell Viability assays in patient-derived GSC-Cas9 lines. Cells were treated with the indicated doses of TMZ for seven days at which point cell viability was assessed via alamarBlue. n=3 biological replicates, mean ± SEM (D) Multicolour competition assay performed in non-neoplastic human neural stem cells with Dox-inducible Cas9 expression. Cells expressing nls-GFP and indicated gRNA were co-cultured with an equal amount of cells expressing nls-mCherry and a non-targeting *(LacZ*) gRNA for a period of 21 days with relative proportion of GFP/mCherry cells calculated every 3-4 days. Depicted are mean ± SEM from n=3 biological replicates (E) Rescue of increased TMZ sensitivity observed in *MCM8^−-/−^* cells upon inducible expression of gRNA-resistant *MCM8* (left). Depicted are mean ± SEM from n=3 biological replicates. *MCM8* expression levels measured using qRT-PCR (right), mean ± SD n=3. (F) Cell Viability assays in patient-derived GSC-Cas9 lines. Cells were treated with the indicated doses of TMZ for seven days at which point cell viability was assessed via alamarBlue, n=3 biological replicates, mean ± SEM. ZC3H7A O.E. represents lentiviral-mediated overexpression of ZC3H7A. (G) BioID of ZC3H7A reveals interacting proteins enriched for stress granule (green), P-Body (red) and nonsense mediated decay (yellow) biological processes. (H) Localization of venus-tagged ZC3H7A in G361NS cells showing localization to granular structures in the cytosol. Data is representative of n=3 replicates, scale, 10 µM. (I) Co-localization of mCherry-G3BP1 and Venus-ZC3H7A in G361NS.

**Table S7:**
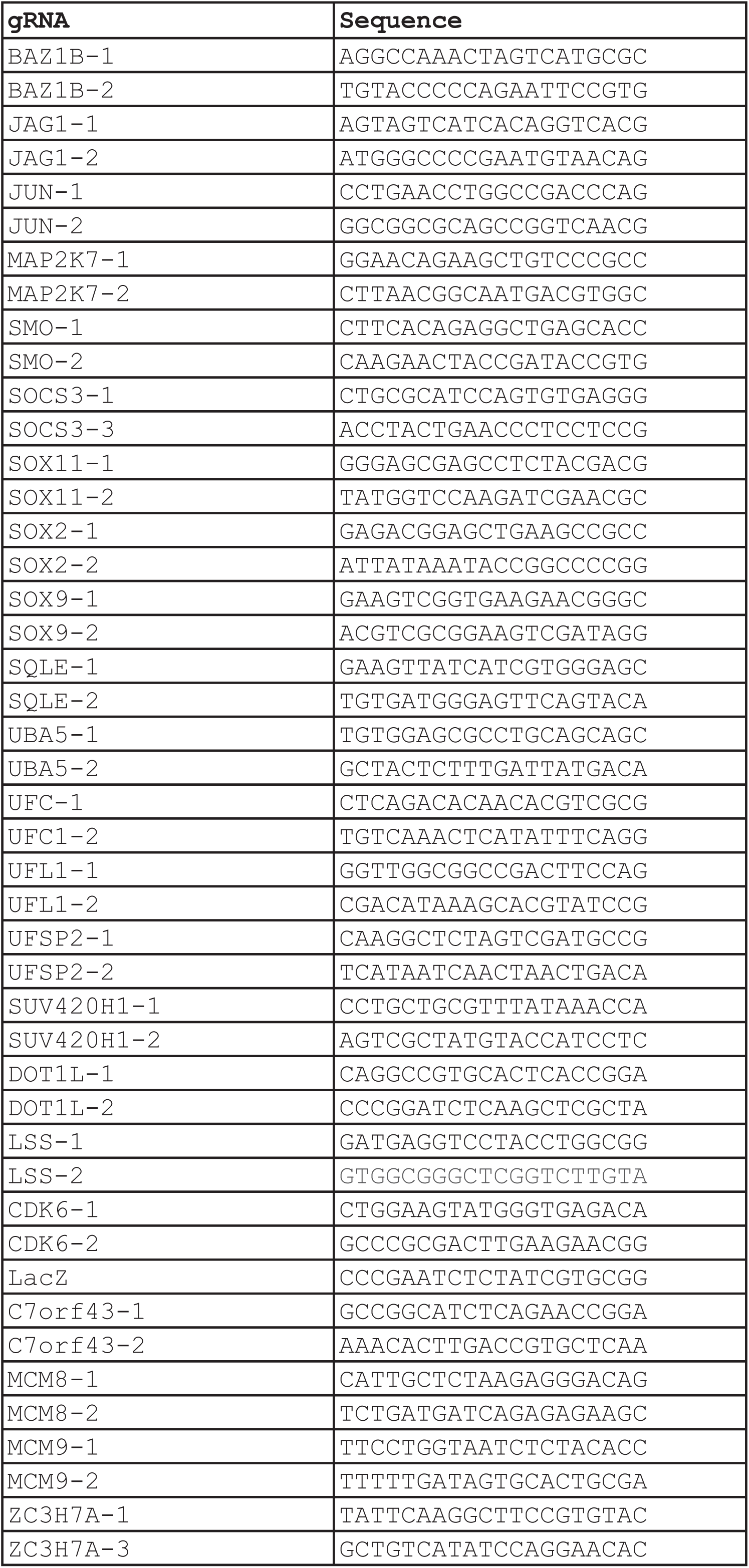
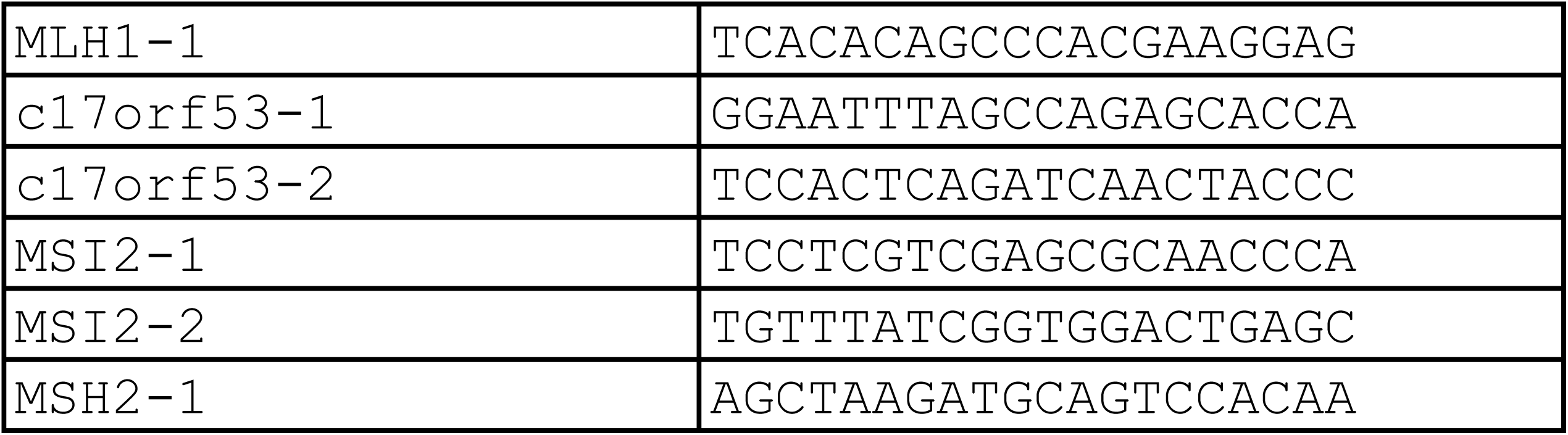
gRNA sequences used in this study. Related to Figures 1,2,5,6,7

**Table S8.**
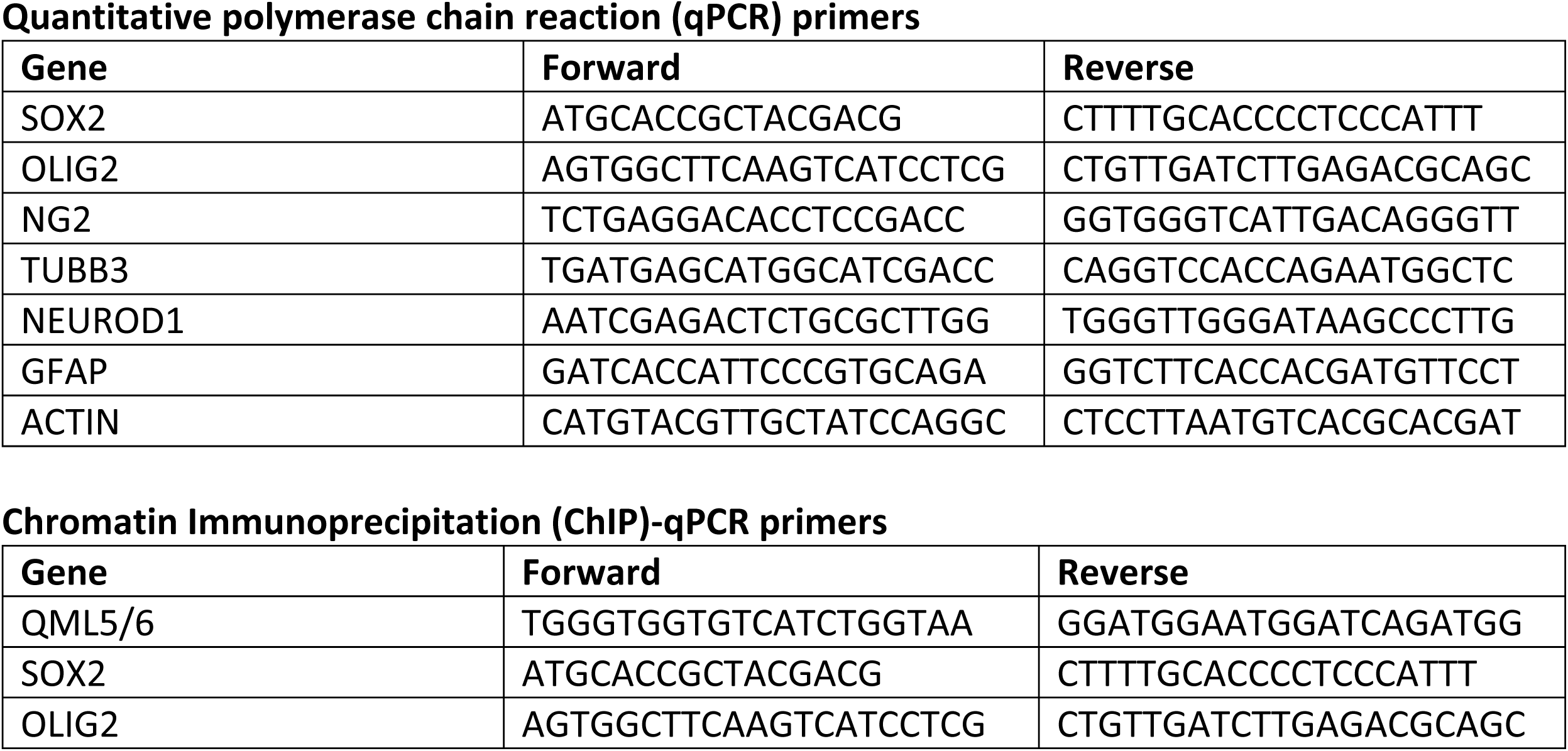
Quantitative polymerase chain reaction (qPCR) primers

## References

Aguirre, A.J., Meyers, R.M., Weir, B.A., Vazquez, F., Zhang, C.-Z., Ben-David, U., Cook, A., Ha, G., Harrington, W.F., Doshi, M.B., et al. (2016). Genomic Copy Number Dictates a Gene-Independent Cell Response to CRISPR/Cas9 Targeting. Cancer Discov. 6, 914–929.

Babicki, S., Arndt, D., Marcu, A., Liang, Y., Grant, J.R., Maciejewski, A., and Wishart, D.S. (2016). Heatmapper: web-enabled heat mapping for all. Nucleic Acids Res. 44, W147–W153.

Bao, S., Wu, Q., McLendon, R.E., Hao, Y., Shi, Q., Hjelmeland, A.B., Dewhirst, M.W., Bigner, D.D., and Rich, J.N. (2006). Glioma stem cells promote radioresistance by preferential activation of the DNA damage response. Nature 444, 756–760.

Ben-David, U., Ha, G., Tseng, Y.-Y., Greenwald, N.F., Oh, C., Shih, J., McFarland, J.M., Wong, B., Boehm, J.S., Beroukhim, R., et al. (2017). Patient-derived xenografts undergo mouse-specific tumor evolution. Nat. Genet. 49, 1567–1575.

Bowman, R.L., Wang, Q., Carro, A., Verhaak, R.G.W., and Squatrito, M. (2017). GlioVis data portal for visualization and analysis of brain tumor expression datasets. Neuro. Oncol. 19, 139–141.

Brennan, C.W., Verhaak, R.G.W., McKenna, A., Campos, B., Noushmehr, H., Salama, S.R., Zheng, S., Chakravarty, D., Sanborn, J.Z., Berman, S.H., et al. (2013). The somatic genomic landscape of glioblastoma. Cell 155, 462–477.

Brinkman, E.K., Chen, T., Amendola, M., and van Steensel, B. (2014). Easy quantitative assessment of genome editing by sequence trace decomposition. Nucleic Acids Res. 42, e168.

Bulstrode, H., Johnstone, E., Marques-Torrejon, M.A., Ferguson, K.M., Bressan, R.B., Blin, C., Grant, V., Gogolok, S., Gangoso, E., Gagrica, S., et al. (2017). Elevated FOXG1 and SOX2 in glioblastoma enforces neural stem cell identity through transcriptional control of cell cycle and epigenetic regulators. Genes Dev. 31, 757–773.

Caiazzo, M., Giannelli, S., Valente, P., Lignani, G., Carissimo, A., Sessa, A., Colasante, G., Bartolomeo, R., Massimino, L., Ferroni, S., et al. (2015). Direct conversion of fibroblasts into functional astrocytes by defined transcription factors. Stem Cell Reports 4, 25–36.

Campeau, E., Ruhl, V.E., Rodier, F., Smith, C.L., Rahmberg, B.L., Fuss, J.O., Campisi, J., Yaswen, P., Cooper, P.K., and Kaufman, P.D. (2009). A versatile viral system for expression and depletion of proteins in mammalian cells. PLoS One 4, e6529.

corp Cancer Genome Atlas Research Network (2008). Comprehensive genomic characterization defines human glioblastoma genes and core pathways. Nature 455, 1061–1068.

Cao, F., Hata, R., Zhu, P., Ma, Y.-J., Tanaka, J., Hanakawa, Y., Hashimoto, K., Niinobe, M., Yoshikawa, K., and Sakanaka, M. (2006). Overexpression of SOCS3 inhibits astrogliogenesis and promotes maintenance of neural stem cells. J. Neurochem. 98, 459–470.

Chen, C.C., Taniguchi, T., and D’Andrea, A. (2007). The Fanconi anemia (FA) pathway confers glioma resistance to DNA alkylating agents. J. Mol. Med. 85, 497–509.

Chen, J., Li, Y., Yu, T.-S., McKay, R.M., Burns, D.K., Kernie, S.G., and Parada, L.F. (2012). A restricted cell population propagates glioblastoma growth after chemotherapy. Nature 488, 522–526.

Choi, H., Larsen, B., Lin, Z.-Y., Breitkreutz, A., Mellacheruvu, D., Fermin, D., Qin, Z.S., Tyers, M., Gingras, A.-C., and Nesvizhskii, A.I. (2010). SAINT: probabilistic scoring of affinity purification–mass spectrometry data. Nat. Methods 8, 70–73.

Coyaud, E., Mis, M., Laurent, E.M.N., Dunham, W.H., Couzens, A.L., Robitaille, M., Gingras, A.-C., Angers, S., and Raught, B. (2015). BioID-based Identification of Skp Cullin F-box (SCF)β-TrCP1/2 E3 Ligase Substrates. Mol. Cell. Proteomics 14, 1781–1795.

Croker, B.A., Kiu, H., and Nicholson, S.E. (2008). SOCS regulation of the JAK/STAT signalling pathway. Semin. Cell Dev. Biol. 19, 414–422.

DeJesus, R., Moretti, F., McAllister, G., Wang, Z., Bergman, P., Liu, S., Frias, E., Alford, J., Reece-Hoyes, J.S., Lindeman, A., et al. (2016). Functional CRISPR screening identifies the ufmylation pathway as a regulator of SQSTM1/p62. Elife 5.

Deshaies, R.J. (2014). Proteotoxic crisis, the ubiquitin-proteasome system, and cancer therapy. BMC Biol. 12, 94.

Dhanasekaran, D.N., and Reddy, E.P. (2008). JNK signaling in apoptosis. Oncogene 27, 6245–6251.

Dolma, S., Selvadurai, H.J., Lan, X., Lee, L., Kushida, M., Voisin, V., Whetstone, H., So, M., Aviv, T., Park, N., et al. (2016). Inhibition of Dopamine Receptor D4 Impedes Autophagic Flux, Proliferation, and Survival of Glioblastoma Stem Cells. Cancer Cell 29, 859–873.

Eden, E., Navon, R., Steinfeld, I., Lipson, D., and Yakhini, Z. (2009). GOrilla: a tool for discovery and visualization of enriched GO terms in ranked gene lists. BMC Bioinformatics 10, 48.

van Galen, P., Kreso, A., Mbong, N., Kent, D.G., Fitzmaurice, T., Chambers, J.E., Xie, S., Laurenti, E., Hermans, K., Eppert, K., et al. (2014). The unfolded protein response governs integrity of the haematopoietic stem-cell pool during stress. Nature 510, 268–272.

Gallo, M., Coutinho, F.J., Vanner, R.J., Gayden, T., Mack, S.C., Murison, A., Remke, M., Li, R., Takayama, N., Desai, K., et al. (2015). MLL5 Orchestrates a Cancer Self-Renewal State by Repressing the Histone Variant H3.3 and Globally Reorganizing Chromatin. Cancer Cell 28, 715–729.

Hart, T., and Moffat, J. (2016). BAGEL: a computational framework for identifying essential genes from pooled library screens. BMC Bioinformatics 17, 164.

Hart, T., Chandrashekhar, M., Aregger, M., Steinhart, Z., Brown, K.R., MacLeod, G., Mis, M., Zimmermann, M., Fradet-Turcotte, A., Sun, S., et al. (2015). High-Resolution CRISPR Screens Reveal Fitness Genes and Genotype-Specific Cancer Liabilities. Cell 163, 1515–1526.

Hart, T., Tong, A.H.Y., Chan, K., Van Leeuwen, J., Seetharaman, A., Aregger, M., Chandrashekhar, M., Hustedt, N., Seth, S., Noonan, A., et al. (2017). Evaluation and Design of Genome-Wide CRISPR/SpCas9 Knockout Screens. G3 7, 2719–2727.

Hsieh, J., and Zhao, X. (2016). Genetics and Epigenetics in Adult Neurogenesis. Cold Spring Harb. Perspect. Biol. 8.

Hu, Y., and Smyth, G.K. (2009). ELDA: extreme limiting dilution analysis for comparing depleted and enriched populations in stem cell and other assays. J. Immunol. Methods 347, 70–78.

Hunter, C., Smith, R., Cahill, D.P., Stephens, P., Stevens, C., Teague, J., Greenman, C., Edkins, S., Bignell, G., Davies, H., et al. (2006). A hypermutation phenotype and somatic MSH6 mutations in recurrent human malignant gliomas after alkylator chemotherapy. Cancer Res. 66, 3987–3991.

Jiang, M., Zhang, W.-W., Liu, P., Yu, W., Liu, T., and Yu, J. (2017). Dysregulation of SOCS-Mediated Negative Feedback of Cytokine Signaling in Carcinogenesis and Its Significance in Cancer Treatment. Front. Immunol. 8, 70.

Jin, X., Kim, L.J.Y., Wu, Q., Wallace, L.C., Prager, B.C., Sanvoranart, T., Gimple, R.C., Wang, X., Mack, S.C., Miller, T.E., et al. (2017). Targeting glioma stem cells through combined BMI1 and EZH2 inhibition. Nat. Med. 23, 1352–1361.

Kim, H., Bhattacharya, A., and Qi, L. (2015). Endoplasmic reticulum quality control in cancer: Friend or foe. Semin. Cancer Biol. 33, 25–33.

Kondo, N., Takahashi, A., Mori, E., Noda, T., Zdzienicka, M.Z., Thompson, L.H., Helleday, T., Suzuki, M., Kinashi, Y., Masunaga, S., et al. (2011). FANCD1/BRCA2 Plays Predominant Role in the Repair of DNA Damage Induced by ACNU or TMZ. PLoS One 6, e19659.

Lan, X., Jörg, D.J., Cavalli, F.M.G., Richards, L.M., Nguyen, L.V., Vanner, R.J., Guilhamon, P., Lee, L., Kushida, M.M., Pellacani, D., et al. (2017). Fate mapping of human glioblastoma reveals an invariant stem cell hierarchy. Nature 549, 227–232.

Lee, J., Kotliarova, S., Kotliarov, Y., Li, A., Su, Q., Donin, N.M., Pastorino, S., Purow, B.W., Christopher, N., Zhang, W., et al. (2006). Tumor stem cells derived from glioblastomas cultured in bFGF and EGF more closely mirror the phenotype and genotype of primary tumors than do serum-cultured cell lines. Cancer Cell 9, 391–403.

Li, W., Xu, H., Xiao, T., Cong, L., Love, M.I., Zhang, F., Irizarry, R.A., Liu, J.S., Brown, M., and Liu, X.S. (2014). MAGeCK enables robust identification of essential genes from genome-scale CRISPR/Cas9 knockout screens. Genome Biol. 15, 554.

Liu, G., Zhang, J., Larsen, B., Stark, C., Breitkreutz, A., Lin, Z.-Y., Breitkreutz, B.-J., Ding, Y., Colwill, K., Pasculescu, A., et al. (2010). ProHits: integrated software for mass spectrometry-based interaction proteomics. Nat. Biotechnol. 28, 1015–1017.

Lu, C., Ward, P.S., Kapoor, G.S., Rohle, D., Turcan, S., Abdel-Wahab, O., Edwards, C.R., Khanin, R., Figueroa, M.E., Melnick, A., et al. (2012). IDH mutation impairs histone demethylation and results in a block to cell differentiation. Nature 483, 474–478.

Martini, M., Pallini, R., Luongo, G., Cenci, T., Lucantoni, C., and Larocca, L.M. (2008). Prognostic relevance of SOCS3 hypermethylation in patients with glioblastoma multiforme. Int. J. Cancer 123, 2955–2960.

Matsuda, K.-I., Sato, A., Okada, M., Shibuya, K., Seino, S., Suzuki, K., Watanabe, E., Narita, Y., Shibui, S., Kayama, T., et al. (2012). Targeting JNK for therapeutic depletion of stem-like glioblastoma cells. Sci. Rep. 2, 516.

Merico, D., Isserlin, R., and Bader, G.D. (2011). Visualizing Gene-Set Enrichment Results Using the Cytoscape Plug-in Enrichment Map. In Methods in Molecular Biology, pp. 257–277.

Meyer, M., Reimand, J., Lan, X., Head, R., Zhu, X., Kushida, M., Bayani, J., Pressey, J.C., Lionel, A.C., Clarke, I.D., et al. (2015). Single cell-derived clonal analysis of human glioblastoma links functional and genomic heterogeneity. Proc. Natl. Acad. Sci. U. S. A. 112, 851–856.

Meyers, R.M., Bryan, J.G., McFarland, J.M., Weir, B.A., Sizemore, A.E., Xu, H., Dharia, N.V., Montgomery, P.G., Cowley, G.S., Pantel, S., et al. (2017). Computational correction of copy number effect improves specificity of CRISPR-Cas9 essentiality screens in cancer cells. Nat. Genet. 49, 1779–1784.

Nishimura, K., Ishiai, M., Horikawa, K., Fukagawa, T., Takata, M., Takisawa, H., and Kanemaki, M.T. (2012). Mcm8 and Mcm9 form a complex that functions in homologous recombination repair induced by DNA interstrand crosslinks. Mol. Cell 47, 511–522.

Parsons, D.W., Jones, S., Zhang, X., Lin, J.C.-H., Leary, R.J., Angenendt, P., Mankoo, P., Carter, H., Siu, I.-M., Gallia, G.L., et al. (2008). An integrated genomic analysis of human glioblastoma multiforme. Science 321, 1807–1812.

Patel, A.P., Tirosh, I., Trombetta, J.J., Shalek, A.K., Gillespie, S.M., Wakimoto, H., Cahill, D.P., Nahed, B.V., Curry, W.T., Martuza, R.L., et al. (2014). Single-cell RNA-seq highlights intratumoral heterogeneity in primary glioblastoma. Science 344, 1396–1401.

Podobinska, M., Szablowska-Gadomska, I., Augustyniak, J., Sandvig, I., Sandvig, A., and Buzanska, L. (2017). Epigenetic Modulation of Stem Cells in Neurodevelopment: The Role of Methylation and Acetylation. Front. Cell. Neurosci. 11, 23.

Pollard, S.M., Yoshikawa, K., Clarke, I.D., Danovi, D., Stricker, S., Russell, R., Bayani, J., Head, R., Lee, M., Bernstein, M., et al. (2009). Glioma stem cell lines expanded in adherent culture have tumor-specific phenotypes and are suitable for chemical and genetic screens. Cell Stem Cell 4, 568–580.

Raizer, J.J., Chandler, J.P., Ferrarese, R., Grimm, S.A., Levy, R.M., Muro, K., Rosenow, J., Helenowski, I., Rademaker, A., Paton, M., et al. (2016). A phase II trial evaluating the effects and intra-tumoral penetration of bortezomib in patients with recurrent malignant gliomas. J. Neurooncol. 129, 139–146.

Reimand, J., Arak, T., Adler, P., Kolberg, L., Reisberg, S., Peterson, H., and Vilo, J. (2016). g:Profiler-a web server for functional interpretation of gene lists (2016 update). Nucleic Acids Res. 44, W83–W89.

Roberts, A.M., Miyamoto, D.K., Huffman, T.R., Bateman, L.A., Ives, A.N., Akopian, D., Heslin, M.J., Contreras, C.M., Rape, M., Skibola, C.F., et al. (2017). Chemoproteomic Screening of Covalent Ligands Reveals UBA5 As a Novel Pancreatic Cancer Target. ACS Chem. Biol. 12, 899–904.

Scott, C.E., Wynn, S.L., Sesay, A., Cruz, C., Cheung, M., Gomez Gaviro, M.-V., Booth, S., Gao, B., Cheah, K.S.E., Lovell-Badge, R., et al. (2010). SOX9 induces and maintains neural stem cells. Nat. Neurosci. 13, 1181–1189.

Shalem, O., Sanjana, N.E., Hartenian, E., Shi, X., Scott, D.A., Mikkelsen, T.S., Heckl, D., Ebert, B.L., Root, D.E., Doench, J.G., et al. (2013). Genome-Scale CRISPR-Cas9 Knockout Screening in Human Cells. Science 343, 84–87.

Shechter, D., Dormann, H.L., David Allis, C., and Hake, S.B. (2007). Extraction, purification and analysis of histones. Nat. Protoc. 2, 1445–1457.

Steinhart, Z., Pavlovic, Z., Chandrashekhar, M., Hart, T., Wang, X., Zhang, X., Robitaille, M., Brown, K.R., Jaksani, S., Overmeer, R., et al. (2017). Genome-wide CRISPR screens reveal a Wnt-FZD5 signaling circuit as a druggable vulnerability of RNF43-mutant pancreatic tumors. Nat. Med. 23, 60–68.

Stupp, R., Mason, W.P., van den Bent, M.J., Weller, M., Fisher, B., Taphoorn, M.J.B., Belanger, K., Brandes, A.A., Marosi, C., Bogdahn, U., et al. (2005). Radiotherapy plus concomitant and adjuvant temozolomide for glioblastoma. N. Engl. J. Med. 352, 987–996.

Sturm, D., Bender, S., Jones, D.T.W., Lichter, P., Grill, J., Becher, O., Hawkins, C., Majewski, J., Jones, C., Costello, J.F., et al. (2014). Paediatric and adult glioblastoma: multiform (epi)genomic culprits emerge. Nat. Rev. Cancer 14, 92–107.

Suvà, M.L., Rheinbay, E., Gillespie, S.M., Patel, A.P., Wakimoto, H., Rabkin, S.D., Riggi, N., Chi, A.S., Cahill, D.P., Nahed, B.V., et al. (2014). Reconstructing and reprogramming the tumor-propagating potential of glioblastoma stem-like cells. Cell 157, 580–594.

Szklarczyk, D., Morris, J.H., Cook, H., Kuhn, M., Wyder, S., Simonovic, M., Santos, A., Doncheva, N.T., Roth, A., Bork, P., et al. (2017). The STRING database in 2017: quality-controlled protein-protein association networks, made broadly accessible. Nucleic Acids Res. 45, D362–D368.

Tang, A.H., and Rando, T.A. (2014). Induction of autophagy supports the bioenergetic demands of quiescent muscle stem cell activation. EMBO J. 33, 2782–2797.

Toledo, C.M., Ding, Y., Hoellerbauer, P., Davis, R.J., Basom, R., Girard, E.J., Lee, E., Corrin, P., Hart, T., Bolouri, H., et al. (2015). Genome-wide CRISPR-Cas9 Screens Reveal Loss of Redundancy between PKMYT1 and WEE1 in Glioblastoma Stem-like Cells. Cell Rep. 13, 2425–2439.

Traver, S., Coulombe, P., Peiffer, I., Hutchins, J.R.A., Kitzmann, M., Latreille, D., and Méchali, M. (2015). MCM9 Is Required for Mammalian DNA Mismatch Repair. Mol. Cell 59, 831–839.

Uhm, J. (2009). Comprehensive genomic characterization defines human glioblastoma genes and core pathways. Yearbook of Neurology and Neurosurgery 2009, 117–118.

Urra, H., Quillien, V., Hetz, C., and Chevet, E. (2017). Endoplasmic reticulum proteostasis in glioblastoma—From molecular mechanisms to therapeutic perspectives. Sci. Signal.

Venteicher, A.S., Tirosh, I., Hebert, C., Yizhak, K., Neftel, C., Filbin, M.G., Hovestadt, V., Escalante, L.E., Shaw, M.L., Rodman, C., et al. (2017). Decoupling genetics, lineages, and microenvironment in IDH-mutant gliomas by single-cell RNA-seq. Science 355.

Wang, J.Y.J., and Edelmann, W. (2006). Mismatch repair proteins as sensors of alkylation DNA damage. Cancer Cell 9, 417–418.

Wang, J., Cazzato, E., Ladewig, E., Frattini, V., Rosenbloom, D.I.S., Zairis, S., Abate, F., Liu, Z., Elliott, O., Shin, Y.-J., et al. (2016). Clonal evolution of glioblastoma under therapy. Nat. Genet. 48, 768–776.

Wang, T., Wei, J.J., Sabatini, D.M., and Lander, E.S. (2013). Genetic Screens in Human Cells Using the CRISPR-Cas9 System. Science 343, 80–84.

Wang, T., Birsoy, K., Hughes, N.W., Krupczak, K.M., Post, Y., Wei, J.J., Lander, E.S., and Sabatini, D.M. (2015). Identification and characterization of essential genes in the human genome. Science 350, 1096–1101.

Wang, Z., Xu, X., Liu, N., Cheng, Y., Jin, W., Zhang, P., Wang, X., Yang, H., Liu, H., and Tu, Y. (2018). SOX9-PDK1 axis is essential for glioma stem cell self-renewal and temozolomide resistance. Oncotarget 9, 192–204.

Waters, N.J., Smith, S.A., Olhava, E.J., Duncan, K.W., Burton, R.D., O’Neill, J., Rodrigue, M.-E., Pollock, R.M., Moyer, M.P., and Chesworth, R. (2016). Metabolism and disposition of the DOT1L inhibitor, pinometostat (EPZ-5676), in rat, dog and human. Cancer Chemother. Pharmacol. 77, 43–62.

Wei, Y., and Xu, X. (2016). UFMylation: A Unique & Fashionable Modification for Life. Genomics Proteomics Bioinformatics IT>14, 140–146.

Wen, P.Y., and Kesari, S. (2008). Malignant gliomas in adults. N. Engl. J. Med. 359, 492–507.

Wiedemeyer, W.R., Dunn, I.F., Quayle, S.N., Zhang, J., Chheda, M.G., Dunn, G.P., Zhuang, L., Rosenbluh, J., Chen, S., Xiao, Y., et al. (2010). Pattern of retinoblastoma pathway inactivation dictates response to CDK4/6 inhibition in GBM. Proc. Natl. Acad. Sci. U. S. A. 107, 11501–11506.

Yoon, C.-H., Kim, M.-J., Kim, R.-K., Lim, E.-J., Choi, K.-S., An, S., Hwang, S.-G., Kang, S.-G., Suh, Y., Park, M.-J., et al. (2012). c-Jun N-terminal kinase has a pivotal role in the maintenance of self-renewal and tumorigenicity in glioma stem-like cells. Oncogene 31, 4655–4666.

Youn, J.-Y., Dunham, W.H., Hong, S.J., Knight, J.D.R., Bashkurov, M., Chen, G.I., Bagci, H., Rathod, B., MacLeod, G., Eng, S.W.M., et al. (2018). High-Density Proximity Mapping Reveals the Subcellular Organization of mRNA-Associated Granules and Bodies. Mol. Cell 69, 517–532.e11.

Zhang, Y., Zhang, M., Wu, J., Lei, G., and Li, H. (2012). Transcriptional regulation of the Ufm1 conjugation system in response to disturbance of the endoplasmic reticulum homeostasis and inhibition of vesicle trafficking. PLoS One 7, e48587.

Zhou, H., Miki, R., Eeva, M., Fike, F.M., Seligson, D., Yang, L., Yoshimura, A., Teitell, M.A., Jamieson, C.A.M., and Cacalano, N.A. (2007). Reciprocal regulation of SOCS 1 and SOCS3 enhances resistance to ionizing radiation in glioblastoma multiforme. Clin. Cancer Res. 13, 2344–2353.

